# Actinobacteria challenge the paradigm: a unique protein architecture for a well-known central metabolic complex

**DOI:** 10.1101/2020.07.24.212647

**Authors:** Eduardo M. Bruch, Pierre Vilela, Norik Lexa-Sapart, Lu Yang, Bertrand Raynal, Pedro M. Alzari, Marco Bellinzoni

## Abstract

α-ketoacid dehydrogenase complexes are large, tripartite enzymatic machineries carrying out key reactions in central metabolism. Extremely conserved across the tree of life, they have so far all considered to be structured around a high molecular weight hollow core, consisting of up to 60 subunits of the acyltransferase component. We provide here evidence that Actinobacteria break the rule by possessing an acetyltranferase component reduced to its minimally active, trimeric unit, characterized by a unique C-terminal helix that affects the oligomerization and the full 3D architecture of the complex. We show that this unique feature is characterized by an insertion, which together with OdhA is found spread over Actinobacteria. This phylum includes organisms or great interest for agriculture, industrial bio-production and many human pathogens as *Mycobacterium tuberculosis*. Moreover, components of this complex are key for *M. tuberculosis* survival in the human host, and its unique core and protein-protein interactions represent potentially “druggable” targets.

## INTRODUCTION

α-oxoacid dehydrogenase complexes constitute a family of three-partite, ubiquitous metabolic complexes devoted to the oxidative decarboxylation of α-ketoacids and the concomitant production of reducing equivalents in form of NADH ^1^. Three such complexes are known: the pyruvate dehydrogenase (PDH), that provides the main entrance of carbon units into the TCA cycle; the 2-oxoglutarate dehydrogenase (ODH), part of the oxidative branch of the TCA cycle; and the branched-chain α-ketoacid dehydrogenase (BCKDH), involved in the catabolism of aliphatic amino acids. These large tripartite complexes share a common molecular architecture organized around a core made by the E2 component, a flexible, multidomain protein which bears the acyltransferase activity required to transfer the acyl group from the decarboxylated substrate to the CoA-SH acceptor; the number of E2 subunits and the symmetry of the core depend on the complex and the species ^1-5^. First shown by the crystal structure of *Azotobacter vinelandii* E2p, the E2 C-terminal catalytic core assumes an obligate homotrimeric state much similar to chloramphenicol acetyltransferase ^6^, with which it also shares the catalytic mechanism ^7^. The observed higher-order oligomerisation states are made possible by intermolecular trimer-trimer interactions mediated by a well conserved, C-terminal 3_10_ hydrophobic helix which makes intermolecular symmetric interactions ^5^, then confirmed on other E2 enzymes and sometimes described as ‘knobs and sockets’ ^3^. These interactions make symmetric, highly oligomeric states which adopt, in most cases, either an octahedral 432 symmetry, eight E2 homotrimers being positioned at the vertexes of a cube, or an icosahedral 532 symmetry, with 20 trimers assembled as a dodecahedron; the number of subunits depends on the complex and the species ^1^. More recently, the presence of an irregularly shaped 42-mer E2 assembly has been described in the archaeon *Thermoplasma acidophilum* ^8^, although this peculiar oligomeric state is still based on the same kind of interactions between the C-terminal helices. Thus, the oligomeric state of the core, responsible for the large size of the complex, was observed in all analyzed complexes and is a trend commonly accepted to be universally conserved in Eubacteria, Archaea and Eukarya. While the reasons for the presence (and evolutionary conservation) of such huge macromolecular scaffolds remain unclear ^9^, active site coupling (transfer of acyl groups between lipoyl domains within the core) has often been proposed as the major advantage ^1,10^. Also, despite the tripartite organization of these complexes as separate E1/E2/E3 enzymes had always been considered as universal, recent work on *Corynebacterium glutamicum* has shown that no separate E2o component (specific to the ODH complex) is present, and that an E2o succinyl transferase domain is fused to E1o, in a protein called OdhA ^11^. The same situation has then been confirmed for the model mycobacterium *M. smegmatis* ^12^. As the lipoyl-binding domain of E2o is absent from the E2o-E1o fusion, the ODH activity depends on functional lipoyl groups provided *in trans*, and proven to be supplied by E2p from the PDH complex ^13^, therefore suggesting the presence of a mixed PDH/ODH supercomplex. By using an integrative structural biology approach, we describe here how *C. glutamicum* E2p, that was expected to serves as the core of the mixed complex, breaks the rule about the oligomeric state of acyltransferase E2 enzymes reducing its size to the minimal, catalytically active trimeric unit. We also provide evidence supporting these features as a common trait of Actinobacteria.

## RESULTS

### *M. tuberculosis* and *C. glutamicum* E2p are both trimeric in solution

To get insight into the oligomeric state of E2p in *Corynebacteriales*, full-length E2p from both *M. tuberculosis* (*Mt*E2p) and *C. glutamicum* (*Cg*E2p) were successfully expressed in *E. coli* and purified to homogeneity. Analytical ultracentrifugation (AUC) was used to assess the oligomeric assembly of the proteins (Table 1). Strikingly, the ratio between the estimated molecular weight in solution and the one predicted from the sequence indicate a trimeric state in solution for both proteins, with no experimental evidence of higher oligomeric states. The same result was observed for *Cg*E2p catalytic domain alone (*Cg*E2p_CD, comprising residues Leu437-Leu675). No larger assembly, suggestive of a 24-mer or 60-mer state, was observed for any of these proteins in the conditions tested (Table 1).

**Table 1:** Oligomerization state of proteins in solution determined by AUC and SAXS

| Protein | locus name | UNIPROT code | Residues | MW monomer from sequence (kDa) | MW estimaton AUC (kDa) | Frictional ratio extrapolated | Number of oligomers (AUC) | MW estimaton SAXS Bayesian (kDa) | Number of oligomers (SAXS) |
| --- | --- | --- | --- | --- | --- | --- | --- | --- | --- |
| *Cg* E2p_FL | cg2421 | Q8NNJ2 | FL | 70,91 | 221 | 2,85 | 3,1 | 243 | 3,4 |
| *Cg* E2p_CD | cg2421 | Q8NNJ2 | 437-675 | 25,40 | 78 | 1,40 | 3,1 | 77 | 3,0 |
| *Mt* E2p_FL | Rv2215 | P9WIS7 | FL | 57,09 | 141 | 1,71 | 2,5 | 318 | 5,6 |
| *Mt* E2b_FL | Rv2495c | O06159 | FL | 41,21 | 731 | 1,63 | 17,7 | 1179 | 28,6 |
| *Mt* E2b_CD | Rv2495c | O06159 | 165-393 | 24,67 | 600 | 1,48 | 24,3 | 479 | 19,4 |

The absence of the expected higher oligomeric structure was further confirmed by size exclusion chromatography coupled to SAXS analysis, were the molecular mass was estimated using MW estimation tools available in ATSAS ^14,15^ (Table 1 and Suppl. Table 1). For both *Cg*E2p_FL and *Cg*E2p_CD the trimeric form is predominant in solution, whereas for *Mt*E2p_FL the analysis shows a molecular weight estimation that could correspond to a dimer of trimers. However, no evidence of a cubic, dodecameric or any other higher oligomeric state was observed for any of the three proteins.

For *Cg*E2p_CD, the chromatogram showed a single peak separated from the void volume fraction. Constant R_g_ value through the peak indicated the presence of a unique oligomeric species. The experimental R_g_ and D_max_ were 2.27 nm and 8.47 nm. The dimensionless Kratky plot shows a gaussian bell shape that decreases to zero consistent with a folded, globular and compact structure, consistently with the AUC results that indicate a trimeric *Cg*E2p_CD protein with low friction coefficient. For the two full-length proteins, *Mt*E2p_FL and *Cg*E2p_FL, the dimensionless Kratky plot shows a non-gaussian bell shape followed by a shoulder that does not immediately decreases to zero, consistent with a properly folded, but not globular, multidomain protein with flexible linkers (Suppl. Fig. 1). This result, together with the high friction coefficient observed by AUC, is suggestive of well folded, highly elongated proteins in both cases.

**Figure 1:**
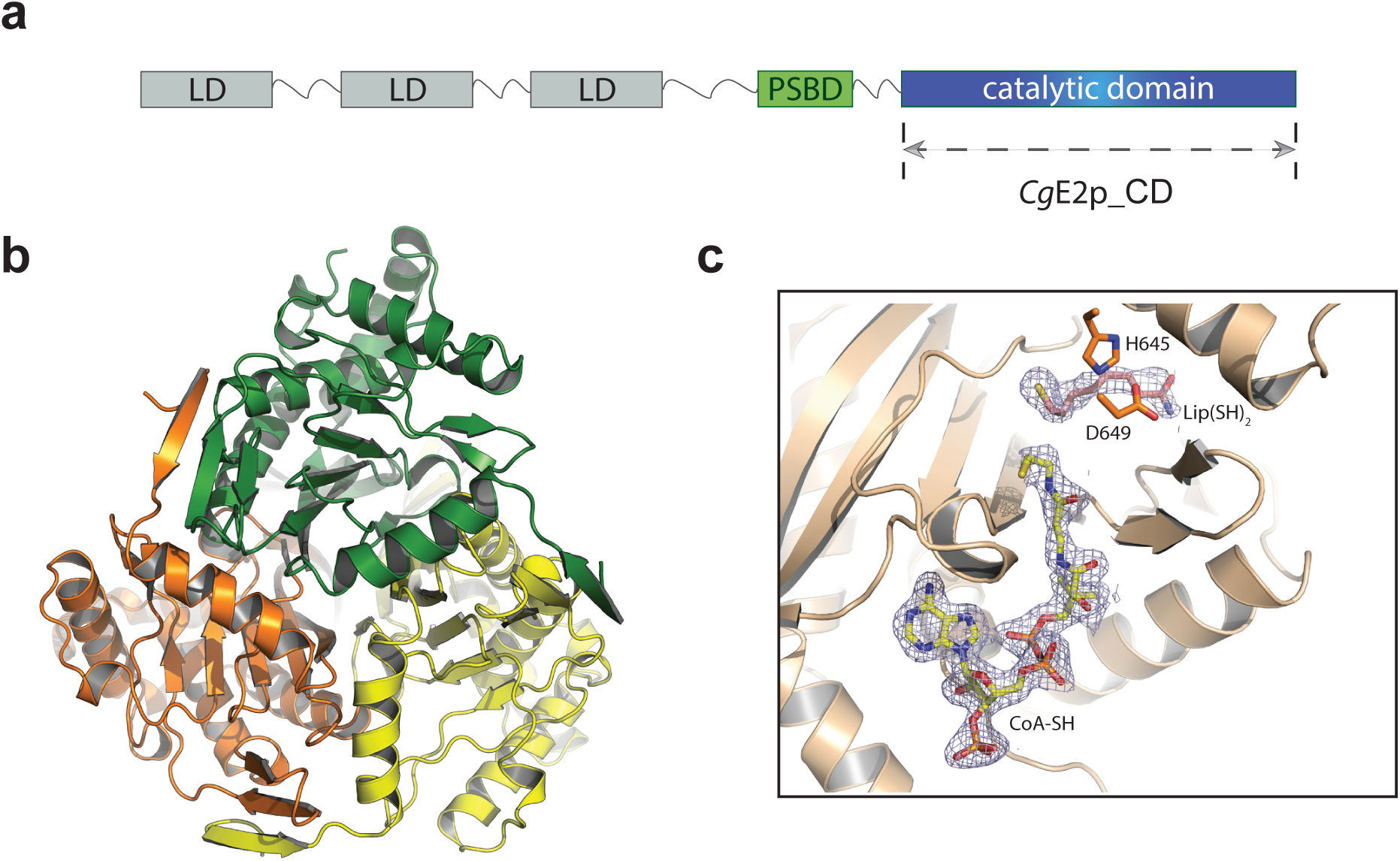
Crystal structure of *Cg*E2p_CD shows a trimeric protein without inter-trimer interactions. **a**- Schematic representation of the domain architechture of *Cg*E2p_FL. The arrow indicates the region selected for the *Cg*E2p_CD construct. **b**- Ribbon representation of the apo form of *Cg*E2p_CD showing its homotrimeric structure, with monomers highlighted in orange, green and yellow, respectively. **c**- Zoomed view of the active site region of ternary complex formed by *Cg*E2p_CD with its sustrates CoA and dihydrolipoamide (Lip(SH)_2_). The 2mFo-DFc Fourier electron density for both ligands, contoured at the 1σ level, is shown as a blue mesh.

### *Cg*E2p_CD posseses a unique C-terminal region characterized by an insertion that blocks the protein in its trimeric form

To understand the reasons for *Cg*E2p behaving as a trimeric protein and not assuming a higher oligomerization state, we decided to characterize it structurally, starting by extensive crystallization trials of *Cg*E2p_CD. Crystals were obtained from the protein in the absence of ligands, in the presence of the acceptor substrate CoA and in a ternary complex with CoA and dihydrolipoamide (Table 2). X-ray diffraction showed that the samples belonged to three different crystallographic space groups, with resolutions ranging from 2.1 to 1.35 Å. In all the crystal forms observed, *Cg*E2p_CD shows the expected, chloramphenicol acetyltransferase-like fold, with an obligate homotrimeric assembly (Fig. 1b), as first observed for the *A. vinelandii* orthologue (*Av*E2p_CD) ^6^. The active site environment at the interface between subunits in the functional trimer is also conserved (Suppl. Fig. 2a), with His645 and Asp649 as the equivalent residues to His610 and Asn614 in *Av*E2p_CD. However, the crystal structure provides no evidence of any higher-order quaternary assembly. To investigate whether the trimeric assembly of *Cg*E2p_CD was still competent to substrate binding, we proceeded to co-crystallization trials with CoA, dihydrolipoamide (Lip(SH)_2_), DTT (as an antioxidant) or their possible combinations. Co-crystallization with CoA yielded high resolution diffracting crystals (Table 2) which show a full-occupancy complex, with no significant conformational change with respect to the *apo* structure. The structure is similar to the equivalent complex from *A. vinelandii* (PDB 1EAD, ^7^), with the adenosine 3’-phosphate moiety occupying basically the same position in both enzymes, CoA assuming a ‘IN’ conformation with the panthetheine chain entering the active site tunnel, and the guanidinium group of Arg480 (Arg450 in *Av*E2p_CD) making a salt bridge with the CoA 3’-phosphate (Suppl. Fig. 2b).

**Table 2.** Crystallographic data collection and refinement statistics (highest resolution shell in parenthesis).

| Data collection | CgE2p_CD | CgE2p_CD<br>Ox-CoA complex | CgE2p_CD<br>CoA/Lip(SH)<sub>2</sub><br>ternary complex | CgE2p_ΔLBDs | CmE2p_CD<br>CoA complex | MtE2b_CD |
| --- | --- | --- | --- | --- | --- | --- |
| Synchrotron<br>beamline | SOLEIL Proxima 2A | SOLEIL Proxima 1 | SOLEIL Proxima 1 | SOLEIL Proxima 1 | ESRF MASSIF-1 | ESRF MASSIF-1 |
| Wavelength (Å) | 0.9801 | 0.9801 | 1.5498 | 0.9786 | 0.9660 | 0.9660 |
| Space group | P3 | P6<sub>3</sub>22 | P2<sub>1</sub>3 | P6<sub>4</sub> | P6<sub>3</sub>22 | F432 |
| Cell dimensions<br>*a*, *b*, *c* (Å) | 158.28, 158.28,<br>63.02 | 73.74, 73.74, 192.19 | 132.13, 132.13,<br>132.13 | 170.39, 170.39, 67.56 | 100.62, 100.62,<br>281.57 | 214.07, 214.07,<br>214.07 |
| $\alpha$, $\beta$, $\gamma$ (°) | 90, 90, 120 | 90, 90, 120 | 90, 90, 90 | 90, 90, 120 | 90, 90, 120 | 90, 90, 90 |
| Resolution (Å) | 79.14 – 1.93<br>(1.96 – 1.93) | 48.05 – 1.35<br>(1.37 – 1.35) | 93.43 – 2.09<br>(2.13 – 2.09) | 85.20 – 2.23<br>(2.37 – 2.23) | 47.38 – 2.50<br>(2.60 – 2.50) | 49.11 – 1.50<br>(1.53 – 1.50) |
| *R*<sub>pim</sub> | 0.043 (0.369) | 0.020 (0.545) | 0.028 (0.334) | 0.027 (0.484) | 0.028 (0.487) | 0.023 (0.391) |
| *I* / $\sigma$(*I*) | 14.0 (2.2) | 16.1 (1.1) | 16.6 (2.2) | 16.0 (1.5) | 18.7 (1.7) | 14.9 (1.9) |
| Completeness (%) | 99.3 (100.0) | 99.6 (92.8) | 100.0 (100.0) | 95.0 (74.8) | 100.0 (100.0) | 100.0 (100.0) |
| CC(1/2) | 0.998 (0.736) | 0.998 (0.844) | 0.999 (0.797) | 0.999 (0.629) | 0.999 (0.593) | 0.999 (0.748) |
| Multiplicity | 3.8 (3.8) | 25.0 (21.2) | 19.2 (19.1) | 20.6 (15.9) | 19.0 (19.6) | 17.2 (17.4) |
| **Refinement** |  |  |  |  |  |  |
| Resolution (Å) | 1.93 | 1.35 | 2.09 | 2.23 | 2.50 | 1.50 |
| No. reflections | 131357 | 68431 | 45719 | 40014 | 30177 | 67274 |
| *R*<sub>work</sub> / *R*<sub>free</sub> (%) | 16.6 / 18.5 | 16.8 / 19.4 | 16.4 / 17.2 | 19.0 / 21.3 | 20.1 / 22.0 | 19.0 / 19.3 |
| No. atoms |  |  |  |  |  |  |
| Protein | 10944 | 1830 | 3659 | 5721 | 3617 | 1668 |
| Ligands/ions | - | 64 | 108 | 17 | 96 | 13 |
| Solvent | 1257 | 300 | 320 | 308 | 131 | 337 |
| Average B-factors |  |  |  |  |  |  |
| Protein | 29.5 | 23.4 | 42.2 | 63.6 | 69.0 | 26.4 |
| Ligand/ions | - | 25.3 | 51.5 | 81.4 | 66.1 | 25.3 |
| Solvent | 38.2 | 38.8 | 51.6 | 58.8 | 64.8 | 40.4 |
| R.m.s deviations |  |  |  |  |  |  |
| Bond lengths (Å) | 0.008 | 0.010 | 0.008 | 0.008 | 0.008 | 0.008 |
| Bond angles (°) | 0.91 | 1.04 | 0.91 | 0.96 | 0.91 | 0.93 |
| PDB entry code | 6ZZI | 6ZZJ | 6ZZK | 6ZZL | 6ZZM | 6ZZN |

**Figure 2:**
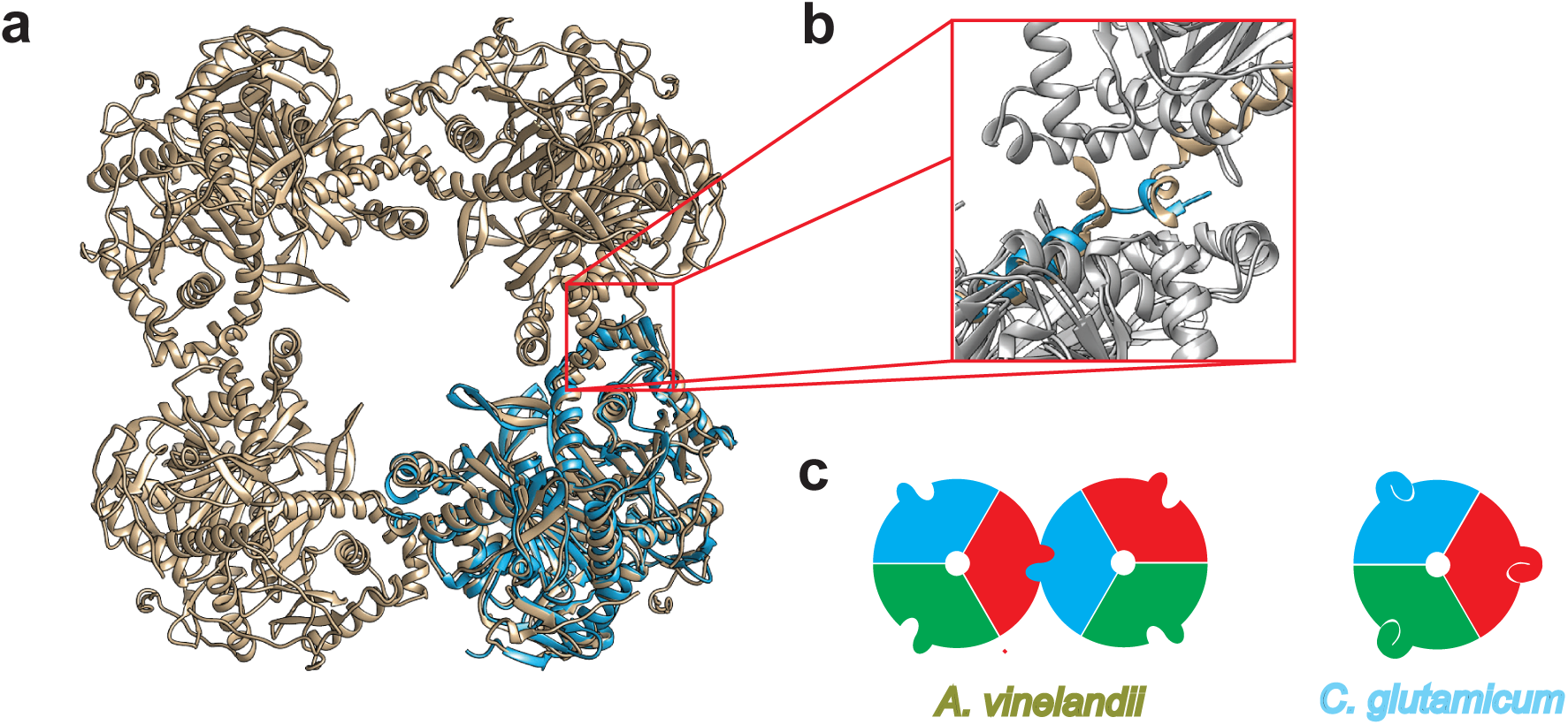
A unique TTI helix orientation is responsable for the absence of inter-trimer interaction. **a**- Ribbon representation of the trimeric *Cg*E2p_CD (cyan) superimposed to *Av*E2p_CD, which forms a 24-mer cubic crystal structure (PDB entry 1EAB). **b**- Zoomed view of the trimer-trimer interaction region showing how the distinct conformation of the TTI helix in *Cg*E2p locates the helix in the position of the incoming TTI helix of *Av*E2p. **c**- On the left, schematic representation of the inter-trimer interaction in *Av*E2p, with different colours for each monomer and the protruding extension representing the C-terminal TTI, allowing the assembly of the 24-mer cube. On the right, the intramolecular interaction of the same TTI helix in *Cg*E2p, hindering trimer-trimer interactions and therefore blocking the protein in a trimeric state.

A ternary complex was obtained by co-crystallization with CoA and free dihydrolipoamide, similarly to *Av*E2p_CD ^7^, although it belonged to a different crystalline form with respect to the CoA complex (Table 2). This structure shows the same, ‘IN’ conformation for the CoA pantetheine chain and the dihydrolipoamide molecule in an orientation close, albeit not identical, to the one observed for the corresponding complex in *Av*E2p_CD (PDB entry 1EAB; Fig. 1c, Suppl. Fig. 3).

**Figure 3:**
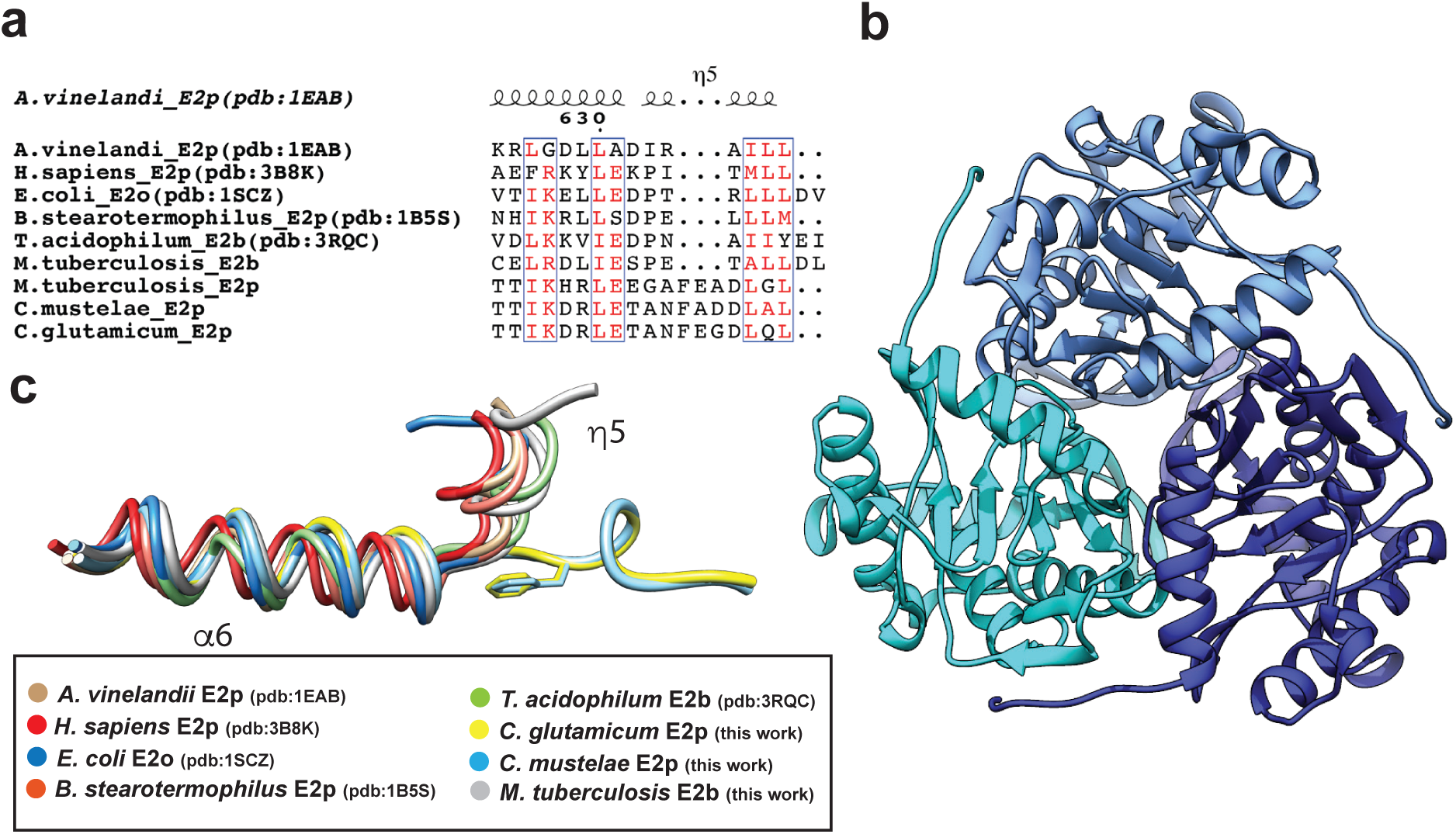
A three residues insertion (PCI) is responsable for the unique TTI helix orientation that blocks inter-trimer interactions. **a**- Sequence alignment, limited to the C-terminal region, between several E2 enzymes of known structure with: *Mt*E2b, *Mt*E2p, *Cg*E2p and *Cm*E2p (this work). **b**- Trimeric structure of *Cm*E2p_CD shown in ribbon representation with different tones of blue per monomer. **c**- Licorized representation corresponding to the C-terminal region of the structural alignment of *Cg*E2p_CD (cyan) and *Cm*E2p_CD (yellow) with: E2p from *A*.*vinelandii, H. sapiens, E. coli, B. stearothemophilus* and *M. smegmatis*, E2o from *E. coli* and E2b from *T. acidophilum*.

Overall, *Cg*E2p_CD is very similar to the available structures of homologous acyltransferase domains from other E2 enzymes, with RMSDs of 1.1-1.3 Å over 220-225 aligned residues (RMSD 1.16 Å over 225 residues), as well as to the other available structures of acyltransferase domains from E2 enzymes, like E2p from *Bacillus stearothermophilus* ^5^ (pdb 1B5S, with which the rms difference is 1.10 Å over 222 residues), E2o from *E. coli* ^16^ (pdb 1SCZ, rmsd 1.29 Å over 211 residues) and E2b from *Bos taurus* ^3^(pdb 2II4, average rmsd 1.3 Å over 221-223 aligned residues), even though the sequence identity is lower than 25% in all cases (Suppl. Fig 5). However, all previously known enzymes share a higher order oligomeric assembly displaying 432 or 532 symmetry, with the functional homotrimers acting as the ‘building blocks’ that interact with each other through their C-terminal, 3_10_ helices ^1,5^. Interestingly, this helix shows the most striking differences when *Cg*E2p_CD is superimposed to *Av*E2p_CD (Fig. 2a/2b and Suppl. Fig 5). While in *Av*E2p_CD this trimer-trimer interacting helix (TTI-helix) is roughly perpendicular to the preceding helix and protrudes outward positioning its hydrophobic residues into a pocket on the facing trimer, in the case of the corynebacterial enzyme the helix lays on the same monomer surface, with the C-terminal hydrophobic residues occupying the same pocket in an intramolecular interaction (Fig. 2b). In other words, the position of the C-terminal helix for *Cg*E2p_CD coincides exactly with the position of the same, incoming helix from the facing trimer in the *A. vinelandii* cubic structure (Fig. 2b). As a consequence, the ‘knobs and sockets’ interaction between trimers is replaced by an intramolecular interaction involving the same features, hindering any trimer-trimer intermolecular interaction and therefore forcing the protein in an homotrimeric state (Fig. 2c).

The above differences are also reflected by the distribution of the hydrophobic residues (Fig. 3a), which in oligomeric *Av*E2p are clustered at the helix C-terminus whereas in *Cg*E2p are arranged at one side of the helix, thus preserving the mostly hydrophobic nature of the helix-pocket interaction. This property is exemplified by the replacement of Leu637 in *Av*E2p for a polar residue (Gln674) in *Cg*E2p, which reflects the new amphipathic nature of the helix. Most importantly, the different orientation of the terminal helix in *Cg*E2p_CD is due to the presence of a three-residue insertion (Phe669-Glu670-Gly671) in *Cg*E2p (Fig. 3a). This insertion allows the C-terminal helix to change its orientation and lay on the surface to reach the accommodating pocket *in cis*, thus replacing an intermolecular interaction by an intramolecular one (Fig. 2c). To confirm that this structural arrangement was not due to the truncated *Cg*E2p_CD protein, a new *Cg*E2p construct including the peripheral subunit binding domain (PSBD), was also overexpressed in *E. coli*, purified and crystallized, yielding a new, 2.5 Å resolution structure (Table 2) showing the catalytic domain in addition to an N-terminal, 13-residue extension that represent part of the flexible linker connecting to the PSBD (Suppl. Fig. 4). Noteworthy, no difference in the oligomeric state, nor in the position of the C-terminal, 3_10_ helix was observed compared to the previous structures. In addition, the E2p ortholog from *M. tuberculosis, Mt*E2p, also possesses a three-residue insertion (Phe547-Glu548-Ala549) in an equivalent position just preceding the predicted 3_10_ terminal helix (Fig. 3a), suggesting that the Phenylalanine-containing insertion (PCI) could be responsible of the lack of higher order oligomerization states.

**Figure 4:**
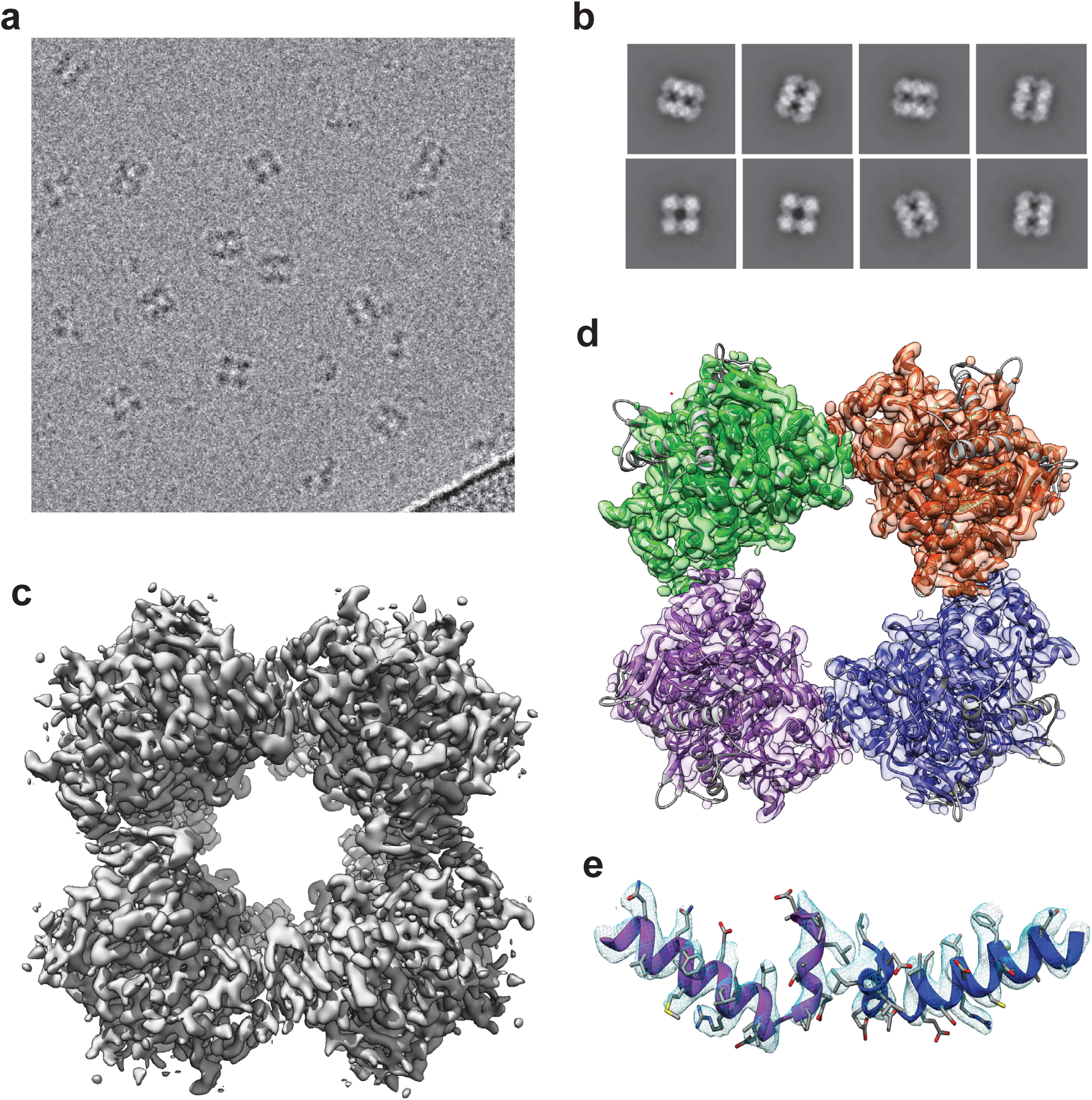
CryoEM structure of *Mt*E2b_CD, an actinobacterial E2b enzyme lacking the PCI, shows canonical TTI helix and quaternary structure assembly. **a**- CryoEM micrograph showing individual particles of *Mt*E2b_CD. **b**- Representative 2D classes showing different particle orientations. **c**- CryoEM map of *Mt*E2b_CD showing its cubic 24-mer with octahedral symmetry. **d**- Crystal structure of the same E2b_CD, extended to the full 24-mer cube (this work), fitted into the EM electron density. **e**- Zoomed region corresponding to the EM electron density of the C-terminal section of the protein, showing the canonical TTI helix orientation observed in other E2 enzymes.

### The Phe containing insertion (PCI) is a common feature in actinobacteria

To analyze the distribution of the PCI, we performed sequence analysis of the E2-coding genes in representative genomes distributed within the phylum Actinobacteria (Suppl. Fig. 6). Our results show that the PCI is not only present in *Corynebacteriales*, including *M. tuberculosis* and *C. glutamicum*, but more generally in sequences from two classes, namely *Actinobacteria* (also called Actinomycetales that represent the biggest class inside the Phylum Actinobacteria) and *Acidimicrobiia* (Suppl. Fig. 6). The high conservation of the Phe residue present in the insertion emphasizes the importance of its interaction within the context of the hydrophobic pocket. Other residues involved in H-bond interactions with the pocket (mainly Glu670, Leu675 and Leu677) are also conserved (Suppl. Fig. 6). Interestingly, although the PCI is conserved, the TTI-helix region has lost some of the C-terminal residues and even the full C-terminal helix is absent in some of the organisms (data not shown), and is in agreement with the absence of the inter-trimer interaction in which this helix is involved.

### PCI and E2 oligomerization

To confirm the relationship between the presence of the three-residue, actinobacterial-specific C-terminal insertion (PCI) and the oligomerization order of E2 proteins from the Actinobacteria class, we followed two parallel paths. First, we structurally analyzed the orthologous protein from *Corynebacterium mustelae*, a recently described species from the same genus ^17^, as an example of orthologue bearing the same insertion (PCI). The catalytic domain of the predicted E2p enzyme from this species (*Cm*E2p_CD, residues 428-666) was crystallized in complex with CoA, and its structure solved by molecular replacement to 2.5 Å resolution (Table 2). The crystal structure shows an homotrimeric arrangement of *Cm*E2p_CD with no significant structural changes with respect to the *C. glutamicum* counterpart (RMSD of 0.79 Å over the whole trimer), and without evidence of any relevant higher order oligomeric state (Fig. 3b). In particular, the C-terminal 3_10_ helix has the same relative orientation as in *Cg*E2p_CD and shows the same intramolecular interaction over the trimer surface (Fig. 3b), with the three-residue insertion (including the well-conserved Phe) occupying an equivalent position in both structures (Fig. 3c and 3a).

Likewise, we analyzed *Mt*E2b, the predicted E2 component of the BCKDH complex (Branched Chain KetoAcid Dehydrogenase) ^18^. The catalytic domain, or *Mt*E2b_CD (residues 165-393), was overexpressed in *E. coli* with an N-terminal, cleavable His_6_ tag and purified following the same experimental strategy. The rationale for the choice of this protein resides in the absence of the PCI, as shown by the sequence alignment in Fig. 3a, and confirmed by the unpublished crystal structure of a similar construct (pdb entry 3L60), issued from a structural genomics initiative. Crystallization experiments carried out after TEV cleavage of the N-terminal His_6_ tag yielded bipyramid shaped crystals belonging to the cubic space group F432 and diffracting X-rays up to 1.5 Å resolution (Table 2). The crystal structure, solved by molecular replacement using the coordinates available from the pdb, shows the 24-mer cubic arrangement observed for *Av*E2p_CD or bovine E2b ^3^ (Suppl. Fig. 8). The terminal 3_10_ helix assumes the same canonical conformation and is involved in the ‘knobs and sockets’ trimer-trimer interaction. However, this is in striking contrast with the pdb model 3L60, which rather shows an homotrimeric arrangement for an analogous construct (residues Pro165-Leu393 as deduced from the information available in the pdb). To verify whether this discrepancy is due to the presence of a C-terminal purification tag leftover in the previously crystallized sample, we carried out further solution studies on both *Mt*E2b_FL and *Mt*E2b_CD. Both AUC data and *ab-initio* models obtained from SAXS data are consistent with a large, cubic assembly for both constructs (Table 1), indicative of the canonical 24-mer assembly (Suppl. Fig. 1 and 9).

Finally, both *Mt*E2b constructs were analyzed by single-particle cryo-EM. Isolated particles indeed show a cubic shape structure for both *Mt*E2b_FL and *Mt*E2b_CD proteins, with a particle side of approximately 13 nm (Fig. 4a and Suppl. Fig. 10). Several rounds of 2D-classification were performed to select a subset of good particles where the class average reflected the expected cube form and high-resolution features such as defined alpha helices (Fig. 4b and Suppl. Fig. 11). Several *ab-initio* models were calculated and, after 3D classification and the exclusion of “junk” particles showing partial dissociation of the complex, a good dataset of particle was selected. After homogeneous refinement, local CTF refinement and density modification the structure was solved and the corresponding cubic assembly was obtained at a resolution of 3.82 Å (Fig. 4c and Suppl. Fig. 11 a and c). Real space refinement was performed to fit the 24-mer, cubic crystallographic assembly into the EM map, showing a very good fitting overall (Fig. 4d) and good density for the TTI helix (Fig. 4e), which, in turn, confirmed its canonical orientation including the ‘knobs and sockets’ interactions, in striking contrast with the E2p situation. The ensemble of our structural and biophysical data therefore supports a strong link between the three-residue C-terminal insertion (PCI), the orientation of the terminal 3_10_ helix (TTI), and the E2 oligomeric state.

### The PCI in actinobacterial E2p correlates with the presence of OdhA

Unlike other canonical E1 proteins, *C. glutamicum* OdhA has a unique domain architecture characterized by the fusion of an E2-like domain to E1o ^13,19^ (Fig. 5d). Taking advantage of this unique E2-E1 fusion feature, an architecture conservation strategy was used to retrieve the sequences of OdhA homologs from an extended list of available genomes. This strategy was based on the search of the pfams associated to both the E2 like domain and E1o (Fig. 5a). This analysis revealed that, except for a few isolated cases, OdhA homologs are a unique feature of Actinobacteria (Fig. 5a). Inside this Phylum, OdhA homologs are widely present in the Actinomycetales class, except for members of the order Bifidobacteriales, and the Acidimicrobiia class (Fig. 5b), but not in the classes Coriobacteriia or Rubrobacteria. In the three members from the class Thermoleophilia a unique architecture is present. In this architecture the first domain (‘2-oxoglutarate_dh_Nterminus’) is replaced by two Lipoyl_biotin domain and a E3-binding domain (Fig. 5d) thus bearing a full fusion of E2 and E1o (called LB-PS-E2-E1). Although speculative at this stage, this protein (never reported before) could perform its activity without the need of a lipoyl domain provided *in trans*, and thus avoiding the interaction with E2. This is coherent with the absence of the N-terminal ‘2-oxoglutarate_dh_Nterminus’, present in OdhA, which is important for the interaction with E2 (Fig. 5d).

**Figure 5:**
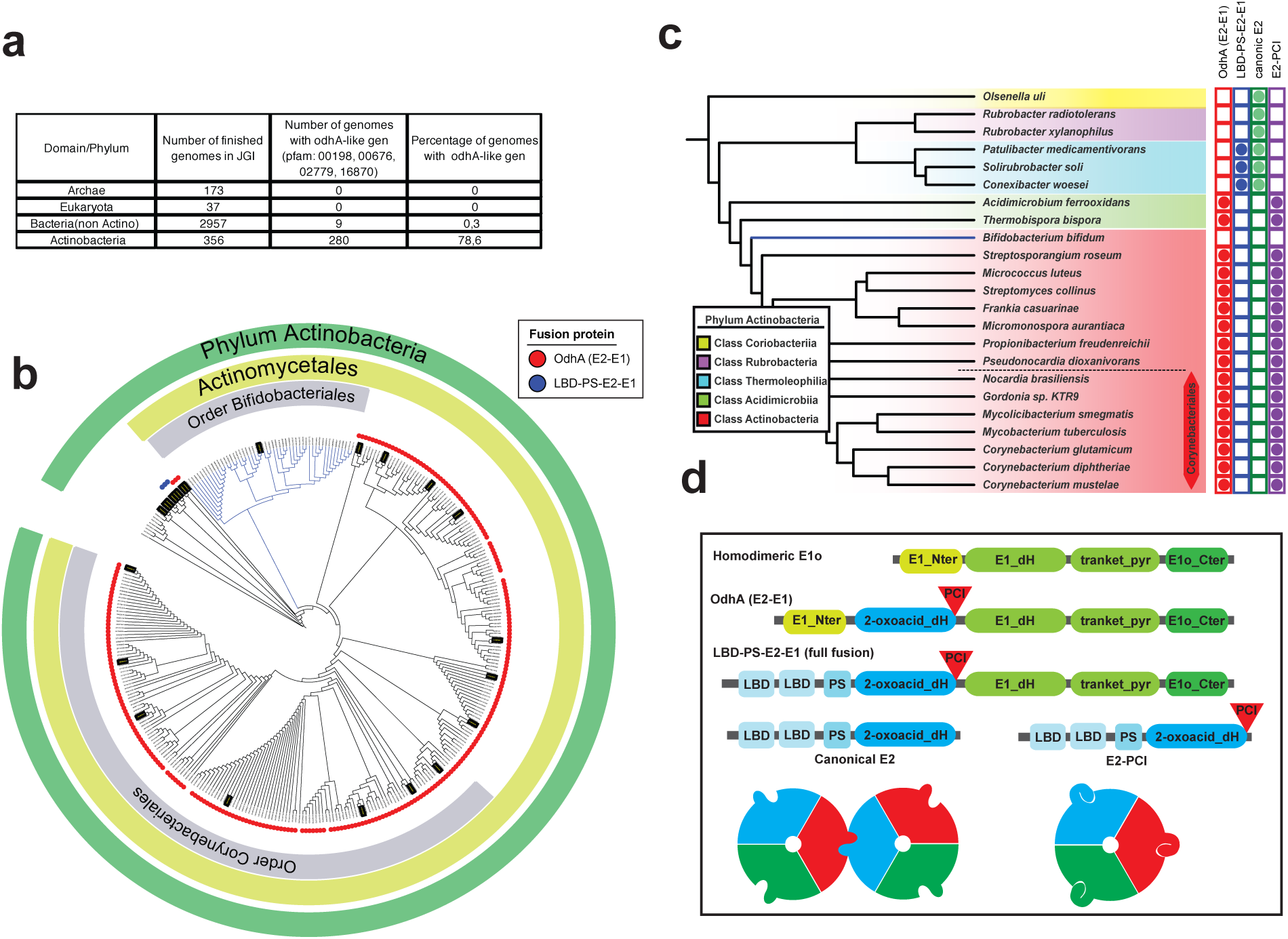
Distribution of OdhA homologs and PCI inside the phylum Actinobacteria. **a**- Table showing the distribution of OdhA homologous found using the conservation of domain architecture predicted from the associated pfams. **b**- Distribution of OdhA homologous (red dots) or the full fusion LBD-PS-E2-E1 (blue dots) inside the phylum Actinobacteria. Branches highlighted in black correspond to genomes selected to be and analyzed for the PCI presence in E2. **c**- Sequences selected from the actinobacterial tree to verify the presence of the PCI.The table shows the correlation between the presence of OdhA and the PCI in E2p. **d**- Domain architechture of E1o, OdhA, the full fusion (LBD-PS-E2-E1) and E2p represented by the pfams used to identify the proteins. Green pfams are associated to homodimeric E1o (Pfam16078:2-oxogl_de-hyd_N, Pfam00676:E1-dh, Pfam02279:Transket_pyr and Pfam16870:OxoGdeHyase_C) while blue pfams are associated to canonical E2 (Pfam00364:Lypoil binding domain, Pfam02817:E3 binding domain and Pfam00198:2-oxoacid dH).

The correlation between the PCI in E2 and the presence of an OdhA homolog (devoid of a lipoyl-binding domain) could be observed for all the representative genomes selected (Fig. 5c). Both features are present in classes Actinobacteria and Acidimicrobiia and absent in the classes Thermoleophilia, Rubrobacteria or Coriobacteria. Since OdhA requires E2p to perform its activity in the context of the PDH/ODH supercomplex, it is tempting to speculate that loss of E2 higher oligomerization could be a feature evolved to optimize the assembly of such a mixed complex, where a E2o-E1o fusion enzyme has to compete with a canonical E1p for the lipoyl groups provided by the same E2p. The latest could explain why this small size E2p was only detected in the actinobacteria phylum and not in any other organism in the tree of life.

## DISCUSSION

The ensemble of enzymatic reactions that make up central metabolism and concur to the production of reducing equivalents in heterotrophic organisms is commonly considered to be extremely conserved. The same has longtime been believed for the respective enzymes and their assemblies, for which most of our current knowledge comes from studies made decades ago on model organisms, which, for prokaryotes, were in most cases represented by *E. coli* or *B. subtilis*. 2-oxoacid dehydrogenase complexes, which are key players in central metabolism, were also object of intensive study during this ‘golden age’ of enzymology, with most of our knowledge coming from seminal work carried out, over several decades, by the groups led by renowned biochemists like Richard N. Perham ^1^, Lester Reed ^20^, and John Guest ^21^. Their extensive biochemical work was then largely confirmed by structural evidence that came out during the ‘90s and the following years, initiated by the work performed in Wim Hol’s lab on the diazotroph, Gram-negative model *A. vinelandii*, and the first determination of a crystal structure of AvE2p_CD ^6^, which in turn confirmed observations dating back to the ‘60s on *E. coli* E2p ^22^ and E2o ^23^. Most of the following structural studies have focused on the PDH complexes, especially the ones from *E. coli, B. stearothermophilus* or their counterparts from human or yeast ^1,24,25^, while ODH and BCKDH complexes, despite a number of biochemical studies ^26^, were largely assumed to share PDH-like molecular structures. In particular, 2-oxoacid dehydrogenase complexes have all been considered to conform to the ‘rule’ of a hollow, large E2 core showing either cubic (432) or icosahedral symmetry (532), assembled according to an ensemble of equivalent, and quasi-equivalent interactions involving the C-terminal end of each E2 monomer ^5^. The more unusual 42-mer E2 assembly described in *T. acidophilum* ^8^, although deviating from the standard symmetry, still possess a hollow core whose assembly is based on the same kind of C-terminal helices interactions, and thus does not represent a real exception to the rule. Questions about the necessity of such large oligomeric assemblies for 2-oxoacid dehydrogenase activity were already raised in the past, when the same team that described the *T. acidophilum* E2 showed that precluding the formation of higher E2 oligomers did not lead to loss of catalytic activity ^9^, in contrast with previous reports. Our work here shows how *Corynebacteriales*, and most *Actinomycetales*, break the ‘dogma’ regarding the three-dimensional architecture of 2-oxoacid dehydrogenase complexes, and raise new questions about the origin and evolution of these enzymatic machineries.

Actinobacteria represent one of the largest and ubiquitous prokaryotic phylum, with more than 50 families. Some of its members are of great economic importance to humans, because agriculture and forests depend on their contributions to the soil system, as *Streptomyces*, one of the largest bacterial genera and a major source of antibiotics. This phylum also includes important human pathogens such as *M. tuberculosis*, as well as major cell factories as *C. glutamicum*. Despite the wide applications of *C. glutamicum* in the biotech industry, which range from the production of amino acids to that of biofuels ^27^, relatively little is known about how *Corynebacteriales* regulate their central metabolism. Since the discovery that the FHA protein OdhI is a phospho-dependent regulator of OdhA, the *C. glutamicum* E1o subunit ^11^, a bunch of evidence indicate that the ODH activity in *Corynebacteriales* – including *M. tuberculosis* and *M. smegmatis* – is regulated by a signal transduction pathway triggered by the Ser/Thr kinase PknG in response to nutrient availability ^28-30^.

The evidence reported here supports a strong correlation between the presence of E2o and E1o activities on a single OdhA-like polypeptide and the loss of highly oligomeric structure of the PDH core, two properties that are most likely related to the dependence of both the PDH and ODH activities on the lipoyl domains provided by a single E2p protein, and, therefore, their coexistence in a mixed multienzymatic machinery. The presence of such kind of supercomplex, first hypothesized in 2006 for *C. glutamicum* ^11^, is in turn suggestive of the presence of regulation mechanisms that might well be unique to Actinobacteria. In this respect, the PknG/GarA (OdhI) signaling pathway, that acts directly on the E1o activity through the OdhI regulator that exerts an allosteric inhibition ^31^, has only be described in Corynebacteriales and might indeed be related to the unusual architecture of the PDH/ODH supercomplex. An additional ODH allosteric regulator is acetyl-CoA, the product of the PDH reaction, identified as an allosteric activator of *M. smegmatis* KGD ^12^ but also, much earlier, as an activator of corynebacterial ODH ^32^, suggesting, in turn, positive feedback mechanisms by product channeling from PDH to ODH. On the other hand, little is known about the regulation of the PDH reaction in Actinobacteria, apart from the known involvement of the RamB transcriptional regulator in the expression of the *aceE* (E1p) gene in *C. glutamicum* ^33^.

Recent developments in metabolomics have started to unravel the mechanisms of metabolic regulation in *M. tuberculosis* ^34,35^, a particularly important facet as metabolic plasticity is a key capability of the pathogen to switch from active replication to dormant states. The strategy of targeting central metabolism for new drugs has been suggested by several groups, mainly due to the presence of unique *M. tuberculosis* pathways that are crucial for pathogen survival and latent state persistence within the host ^34,36,37^. Unlike other central metabolic enzymes, the low similarity with its eukaryotic counterparts makes of the PDH/ODH supercomplex a good drug target candidate. Indeed, the enzymatic components of the complex were shown to participate in the pathogen response to oxidative stress ^38^ or virulence ^18^. In particular, *M. tuberculosis* DlaT (*Mt*E2p), that we show here as being structured as an homotrimer, has been shown to be required for TB infection in the Guinea pig model, and was also involved in the pathogen’s response to reactive nitrogen intermediates ^36^. Because of its unique quaternary arrangement, the components of the PDH/ODH supercomplex are good candidates to reveal new ‘druggable’ surfaces, allowing to target essential, but previously unknown, protein-protein interactions. By revealing the structural principles of the atypical molecular assembly of these enzymes, our work thus opens the way to the development of novel therapeutic approaches.

## MATERIAL AND METHODS

### Plasmid construction

*C. glutamicum* open reading frames were amplified from genomic DNA by PCR and cloned in pET-28a vector by restriction free cloning ^39^, while expression constructs for the other proteins (*Cm*E2p, *Mt*E2p, *Mt*E2b) were provided by Genscript (Leiden, the Netherlands). In all cases, a sequence coding for a His_6_-tag followed by the TEV protease cleavage site (ENLYFQG) were fused to the N-terminus of the protein of interest. Constructs were made to express: for *Cg*E2p (Uniprot accession no. Q8NNJ2), the full-length protein (*Cg*E2p_FL, residues 1-675), the catalytic domain (*Cg*E2p_CD, corresponding to residues 437-675), and the catalytic core and the peripheral subunit binding domain (PSBD; *Cg*E2p-ΔLBDs, corresponding to residues 372 to 675). For *Cm*E2p (Uniprot accession no. A0A0G3H170), the catalytic core was expressed (*Cm*E2p_CD, residues 428-666), while for *Mt*E2p (Uniprot accession no. P9WIS7), the full-length protein was expressed (*Mt*E2p_FL, residues 1-553).

For *Mt*E2b (Uniprot accession no. O06159), two constructs were built: the first one corresponding to the full-length protein (*Mt*E2b_FL, residues 1-393), and the second one corresponding to the catalytic core region (*Mt*E2b_CD, defined as residue 165-393). All plasmids were verified by DNA sequencing.

### Protein purification

Each plasmid was transformed into *E. coli* BL21DE3 using standard protocols. In the case of *Cg*E2p_FL and *Cg*E2p_CD, transformed cells were growth in LB medium at 37 °C until optical density measure at 600 nm reach ∼0.6. Afterwards, temperature was dropped to either 18 °C or 30 °C (see table below) and incubated for 18 hours after adding 0.5 mM isopropyl-β-D-thiogalactopyranoside (IPTG). For *Mt*E2p_FL, *Mt*E2b_FL or *Mt*E2b_CD autoinduction buffer was used so cells were growth at 37 °C for 4 hours and then temperature was dropped to either 18 or 30 °C and cells incubated overnight.

Cells were recovered by centrifugation and washed twice with the corresponding Lysis buffer (see buffer table). Cell lysis was performed using a CF2 cell disruptor (Constant Systems Ltd., Daventry, UK). The soluble fraction was separated from debris by centrifugation at 26,800 g for 1 hour. The recombinant proteins were purified by Ni^2+-^IMAC on HisTrap columns (GE Healthcare). Fractions, as confirmed by SDS-PAGE, containing the protein of interest were dialyzed against GF buffer (see buffer table), supplemented with 1 mM dithiothreitol (DTT), for 12 hours at 18°C, after adding His-tagged TEV protease ^40^ at a 1:30 ratio (w/w) to remove the N-terminal His_6_ tag from the recombinant protein. The cleaved, recombinant proteins were then separated from the His_-_tagged TEV protease (and the cut N-terminal portion) by gravity-flow separation on Ni-NTA agarose (Qiagen). Finally, the recovered recombinant proteins were further purified by size exclusion chromatography using a Sephacryl S-400 16/60 column (GE Healthcare). Fractions containing the protein were pulled, concentrated and flash frozen in liquid nitrogen.

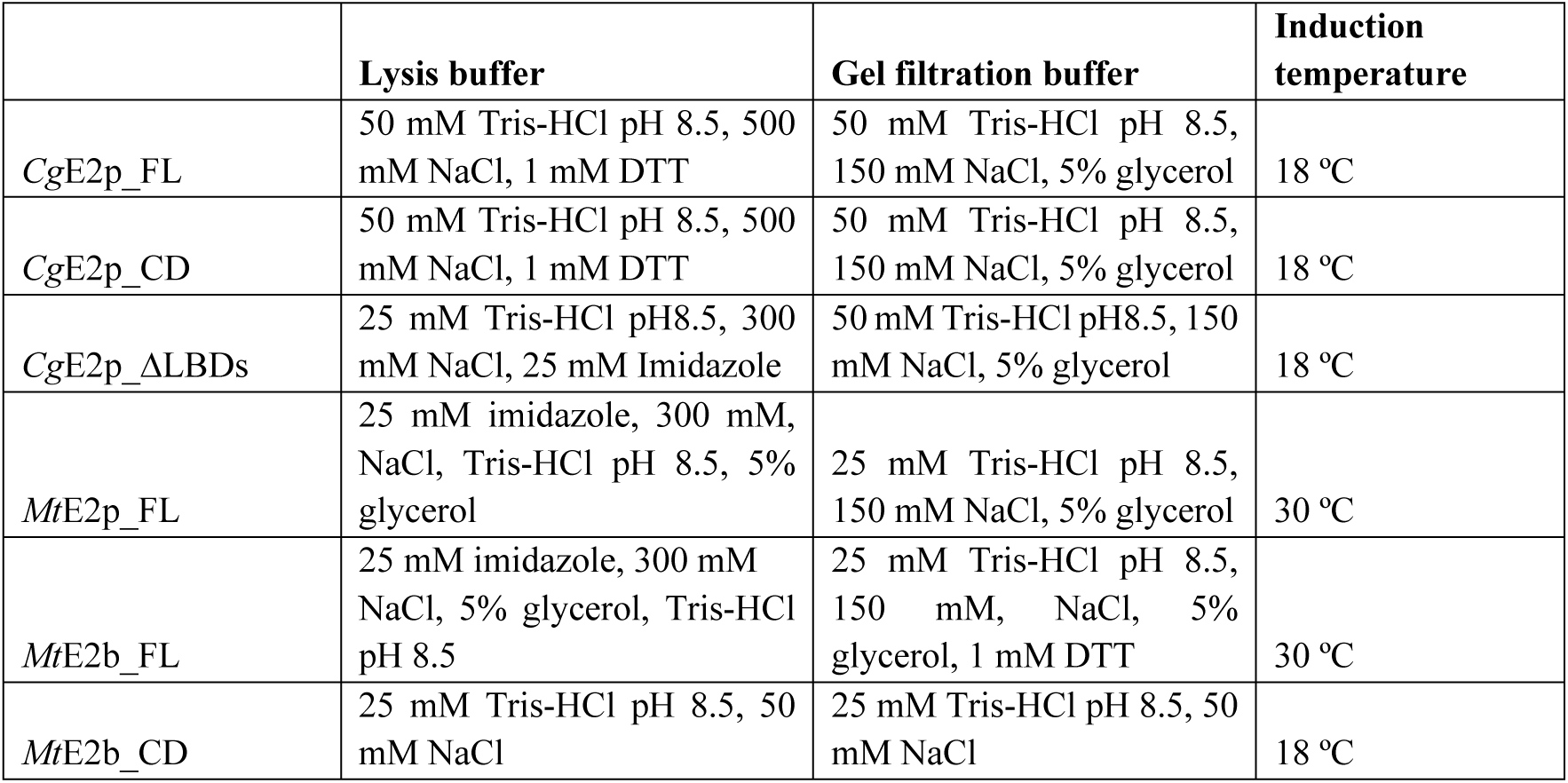

### SEC-SAXS and ab initio modelling

SAXS experiments were performed at SOLEIL synchrotron on the SWING beamline. The beam wavelength was set to 1.033 Å. The detector was located at a distance of 2 meters to collect data at q-range between 0.005 and 0.55 Å^-1^ (q=4π sin θ/λ and 2θ is the scattering angle). Protein sample were injected into a size exclusion column (Superose™ 6 Increase 5/150 GL, GE Healthcare Bio-Sciences) previously equilibrated with corresponding buffer and eluted directly into the SAXS flow-through capillary cell at a flow rate of 0.2 ml/min. Frames were collected continuously during the whole protein elution time.

Frames were analyzed using FOXTROT software. Buffer corresponding frames were averaged and subtracted from sample frames. The forward scattering I(0) and the Rg for each sample subtracted frame were derived from Guinier approximation. The I(0) vs frame curve was used to select the region corresponding to the protein’s elution peak. Frames corresponding to the elution peak and with constant Rg were averaged. All subsequent data processing was performed using ATSAS suite ^14^.

PRIMUS software, from the ATSAS suite, was used to recalculate the I(0) and Rg, while GNOM was used to calculate the pair-distance distribution function P(r) and Dmax ^15^. Finally, the POROD volume and the tools available in ATSAS were used to estimate the molecular weights of the proteins ^41^. The dimensionless Kratky plot was calculated according to previous work ^42,43^, using the value determined by Guinier analysis.

*Ab initio* modelling was also performed using software from the ATSAS suite. For *Mt*E2b_CD, 10 independent models were generated using DAMMIF, with no imposed symmetry. In both cases best models were obtained assuming oblate anisometry. The models were then compared and align with DAMSEL and DAMSUP and output files were generated using DAMAVER, DAMFILT and DAMSTART, all within the ATSAS suite. Finally, the resulting damstart file was used in DAMMIN to obtain a dummy atom model that best fit the data.

### AUC

Sedimentation velocity (SV) analytical ultracentrifugation assays were performed using a Beckman Coulter ProteomeLab XL-I analyticalultracentrifuge equipped with UV-Vis absorbance and Raleigh interference detection systems, using the 8-hole Beckman An-50Ti rotor at 20°C. The speeds chosen are 10,000 rpm for *Mt*E2b_CD and *Mt*E2b_FL and 36,000 rpm for *Mt*E2p_FL, *Cg*E2p_FL and *Cg*E2p_CD. Three concentrations of each protein were prepared for this experiment in their corresponding buffer and loaded into analytical ultracentrifugation cells. During the run SV was followed using by measuring absorbance at 290 nm.

SEDFIT 15.01 (available at http://analyticalultracentrifugation.com; ^44^) was used to calculate the sedimentation coefficient distribution C(s), then corrected to standard conditions to get the final standard values. These coefficients were plotted as a function of the different concentrations and an extrapolation to zero concentration was made to obtain the standard value for the main oligomer. From these values, molecular mass and friction ratio were obtained.

### Crystallization

The crystallization experiments were performed at either 18°C or 4 °C by the sitting drop vapor diffusion technique in 96-well plates, according to established protocols at the Crystallography Core Facility of the Institut Pasteur ^45^. Crystallization conditions were: for *Cg*E2p_CD in apo form, 0.1 M Hepes-Na pH 7.5, 5% PEG4000, 30% 2-methyl-2,4-penthane diol (MPD), at 18°C; for *Cg*E2p_CD in complex with CoA, 0.1 M Hepes-Na pH 7.5, 1.56 M tri-sodium citrate, at 4°C; for *Cg*E2p_CD in ternary complex with CoA and dihydrolipoamide, 0.1 M Tris-HCl pH 7.3, 1.62 M tri-sodium citrate, at 18°C; for *Cg*E2p_ΔLBDs, 30% PEG1500 at 18°C; for *Cm*E2p_CD in complex with CoA, 0.1 M Hepes-Na pH 7.5, 150 mM NaCl, 30% PEG4000, at 18°C; for *Mt*E2b_CD, see 0.1M Imidazole pH 6.5, 4 M NH_4_^+^ acetate at 18°C.

### Data collection and structure solution

Diffraction data were acquired from crystals maintained at 100 K, either at the SOLEIL Synchrotron (Saint-Aubin, France), on the beamline Proxima-1 or Proxima-2A, or on the automated beamline MASSIF-1 at the ESRF (Grenoble France) ^46^. The data were processed with XDS ^47^, run through either XDSME (available at https://github.com/legrandp/xdsme) or autoPROC ^48^, and scaled with Aimless, available from the CCP4 suite ^49^, or STARANISO as provided within autoPROC. The structures were solved by molecular replacement through the program PHASER ^50^. Coordinates of the *E. coli* E2p catalytic domain (PDB entry 4N72; ^51^) were used as the search model to first solve the structure of *Cg*E2p_CD in apo form, which, in turn, served as molecular replacement search model for the following datasets, including *Cm*E2p_CD, but not MtE2b_CD whose structure was solved by using the previously released coordinates of the same protein (pdb entry 3L60). All rebuilding and adjustments of the models were performed with COOT ^52^. The refinement was carried out with BUSTER, applying local structure similarity restraints for non-crystallography symmetry (NCS) ^53^ and a Translation-Libration-Screw (TLS) model. Chemical dictionaries for ligands were generated with the Grade server (http://grade.globalphasing.org). Validation of models was performed with MOLPROBITY ^54^ and the validation tools in the PHENIX suite ^55^. Data collection, refinement, model statistics and pdb accession codes for coordinates and structure factors are indicated in Table 2. Graphical representations were rendered with Pymol ^56^.

### CryoEM

For grid preparation 3.5 µl of 0,2 mg/ml *Mt*E2b_CD were applied into a Lacey Carbon Film on 200 Mesh Copper Grids. Grid vitrification was performed in a Vitrobot (FEI) with 95% humidity, ashless filter paper (Standard Vitrobot Filter Paper, Ø55/20mm, Grade 595) and using a blot time of 3.5 seconds and a blotting force of 0. Grids were then storage and collected in a 200 keV Thermo Scientific Glacios^®^ Cryo-Transmission Electron Microscope. Images were recorded using a Falcon IIIEC direct electron detector in linear mode. The images were collected using EPU software 2.2.0.65REL, with an under focus range from 0.8 μm to 2.6 μm. The pixel size was set to 0.96 Å. Per movie 37 frames were collected in total, with an overall dose of 42.5 e-/Å^2^.

Movies frames were aligned using Motiocor2 ^57,58^, and CTF estimation was performed using Gctf ^59^. Movies were imported and analyzed in Cryosparc v2.15 ^60^. Using the curate exposure feature, 1235 out of 1476 movies, followed by 2483 out of 3533 movies from a second data collection were selected for further analysis. A first round of manual picking was performed and after two rounds of 2D classification, the three most populated classes were selected for template-based particle picking against a dataset containing the selected micrographs. The selected particles were cleaned using the inspect pick function of Cryosparc and three rounds of 2D classification and a final dataset of 40805 and 168705 good particles, corresponding to collection session 1 and 2, respectively, was selected.

A round of *ab initio* and 3D classification into 4 classes was performed to select a subset of 111891 particles that could be refined to 5.8 Å. After local CTF refinement and imposing Octahedral O3 symmetry, this dataset could be refined to 3.92 Å. Finally, density modification was applied using the tool resolve_cryo_em from the PHENIIX suite ^61^, reaching a final map at 3.82 Å resolution. Model fitting into cryo-EM maps was performed using the programs UCSF Chimera ^62^ and phenix.real_space_refine ^63^. The final, sharpened map from Cryosparc as well as the density modified map from resolve_cryo_em were deposited to the EMDB under the accession code 11600. Figures were generated and rendered with UCSF Chimera.

### Bioinformatic analysis

#### a0) Analysis of PCI insertion

Multiple sequence alignments were performed using MUSCLE (https://www.ebi.ac.uk/Tools/msa/muscle/) ^64^. Further alignment details were carried out in AliView version 1.26 ^65^. Sequences from *Mt*E2b, *Mt*E2p, *Cg*E2p, *Cm*E2p and E2 proteins with known structure were compare to the representative sequences to account for the PCI presence. The protein sequence from E2 proteins with known structure were obtained from the pdb database (pdb entries 1EAB, 3B8K, 1SCZ, 3RQC, 1B5S). The final multiple sequence alignment was merged with the pdb entry 1EAB to obtain final figures using ESPript 3.0 (http://espript.ibcp.fr) ^66^. Finally, the presence of the PCI was analyzed in the region corresponding to the C-terminal α-helix (residues 630 to 637 in *Av*E2p).

#### a1) OdhA like gene distribution inside selected genomes

To identify the OdhA like genes into the genomes the gene search tool from IMG/M was used (Integrated Microbial Genomes & Microbiomes system v.5.0) ^67^. To simplify the analysis, only genomes annotated as finished were selected. For each genome OdhA like genes were identified by their conserved Pfam architecture using the associated Pfams: Pfam16078 (2-oxogl_dehyd_N), Pfam00198 (2-oxoacid dH), Pfam00676 (E1-dh), Pfam02279 (Transket_pyr) and Pfam16870 (OxoGdeHyase_C).

| Domain/Phylum | Number of finished genomes in JGI | Number of genomes with *odhA*-like genes (pfam: 16078, 00198, 00676, 02779, 16870) | Percentage of genomes with *odhA*-like genes |
| --- | --- | --- | --- |
| Archae | 173 | 0 | 0 |
| Eukaryota | 37 | 0 | 0 |
| Bacteria (non Actino) | 2957 | 9 | 0,3% |
| Actinobacteria | 356 | 280 | 78,6% |

#### b) odhA like gene distribution inside Phylum Actinobacteria

All the finished genomes corresponding to the phylum actinobacteria and two non-finished genomes corresponding to the class Thermoleophila were selected from IMG/M available genomes (Thermoleophila counts to the date with only one finished genome). After elimination of duplicated genomes OdhA like genes and E2-E1 like genes were identified using its corresponding Pfam architecture.

For E2-E1 like genes the associated genomes correspond to: Pfam00364 (Lypoil binding domain), Pfam02817 (E3 binding domain), Pfam00198 (2-oxoacid dH), Pfam00676 (E1-dh), Pfam02279 (Transket_pyr) and Pfam16870 (OxoGdeHyase_C). For the selected genomes a tree was created using the JGI distance tree tool. The tree is created using the alignment of 16S genes based on the SILVA database and dnadist and neighbor tools from the PHYLIP package (available at https://evolution.genetics.washington.edu/phylip/). Figures were built using the Interactive Tree Of Life (iTOL) v4 online tool ^68^.

#### c) Correlation between odhA like genes and the presence of the PCI in E2p

A representative pool of genomes from the larger orders inside actinobacteria class (namely Corynebacteriales, Streptomycetales, Propionibacteriales, Micromonosporales, Microccocales and Bifidobacteriales) and representatives from the other classes inside the Phylum actinobacteria were selected. For each genome, the corresponding E2 sequences were identified using its corresponding Pfam architecture, namely Pfam00364 (Lypoil binding domain), Pfam02817 (E3 binding domain), Pfam00198 (2-oxoacid dH). For each genome sequence alignments were performed as previously described to assess the existence of the PCI (as described previously).

## ACKNOWLEDGMENTS

This work was funded by the ANR projects SUPERCPLX (ANR-13-JSV8-0003) and METACTINO (ANR-18-CE92-0003), both granted to M.B., and by institutional grants from the Institut Pasteur and the CNRS. We are grateful to Ahmed Haouz, Patrick Weber and Cédric Pissis (Crystallography Core Facility, Institut Pasteur) for carrying out robot-driven crystallization screenings. We also acknowledge Sébastien Brûlé (Molecular Biophysics Core Facility) for his assistance in AUC experiments, and Francesca Gubellini (Structural Microbiology) for her help in the sample preparation for negative staining EM grids. The NanoImaging Core at Institut Pasteur is acknowledged for support with sample preparation, image acquisition and analysis, and its whole staff (Jean-Marie Winter, Stéphane Tachon and Matthijn Vos) is gratefully acknowledged for the supportive help. The NanoImaging Core was created with the help of a grant from the French Government’s ‘Investissements d’Avenir’ program (EQUIPEX CACSICE – “Centre d’analyse de systèmes complexes dans les environnements complexes”, ANR-11-EQPX-0008). L.Y. was funded by CSC (China Scholarship Council) and is part of the Pasteur - Paris University (PPU) International PhD program. The PPU project has received funding from the CNBG (China National Biotec Group Company Limited). We also acknowledge the synchrotron sources Soleil (Saint-Aubin, France), and ESRF (Grenoble, France) for granting access to their facilities, and their staff for helpful assistance on the respective beamlines. We would also like to thank Juan José Pierella Karlusich (ENS, Paris) and Aurélien Thureau (Synchrotron Soleil) for many helpful discussions.

## AUTHOR CONTRIBUTIONS

E.M.B., P.V., N.L.-S. and L.Y. generated expression plasmids and produced recombinant proteins; E.M.B., P.V., L.Y. and M.B. collected X-ray diffraction data; M.B. solved and refined crystallographic structures; E.M.B., P.V., N.L-S. and B.R. collected and analyzed AUC data; E.M.B. and B.R. collected and analyzed SAXS data; E.M.B. collected and analyzed cryo-EM data, and performed phylogeny analysis; P.M.A. provided advice on data interpretation and contributed to the manuscript; M.B. supervised the work; E.M.B. and M.B. wrote the paper. All authors reviewed the manuscript and agreed on its content.

