## Supplementary figures&tables for "Actinobacteria challenge the paradigm: a unique protein architecture for a well-known central metabolic complex"

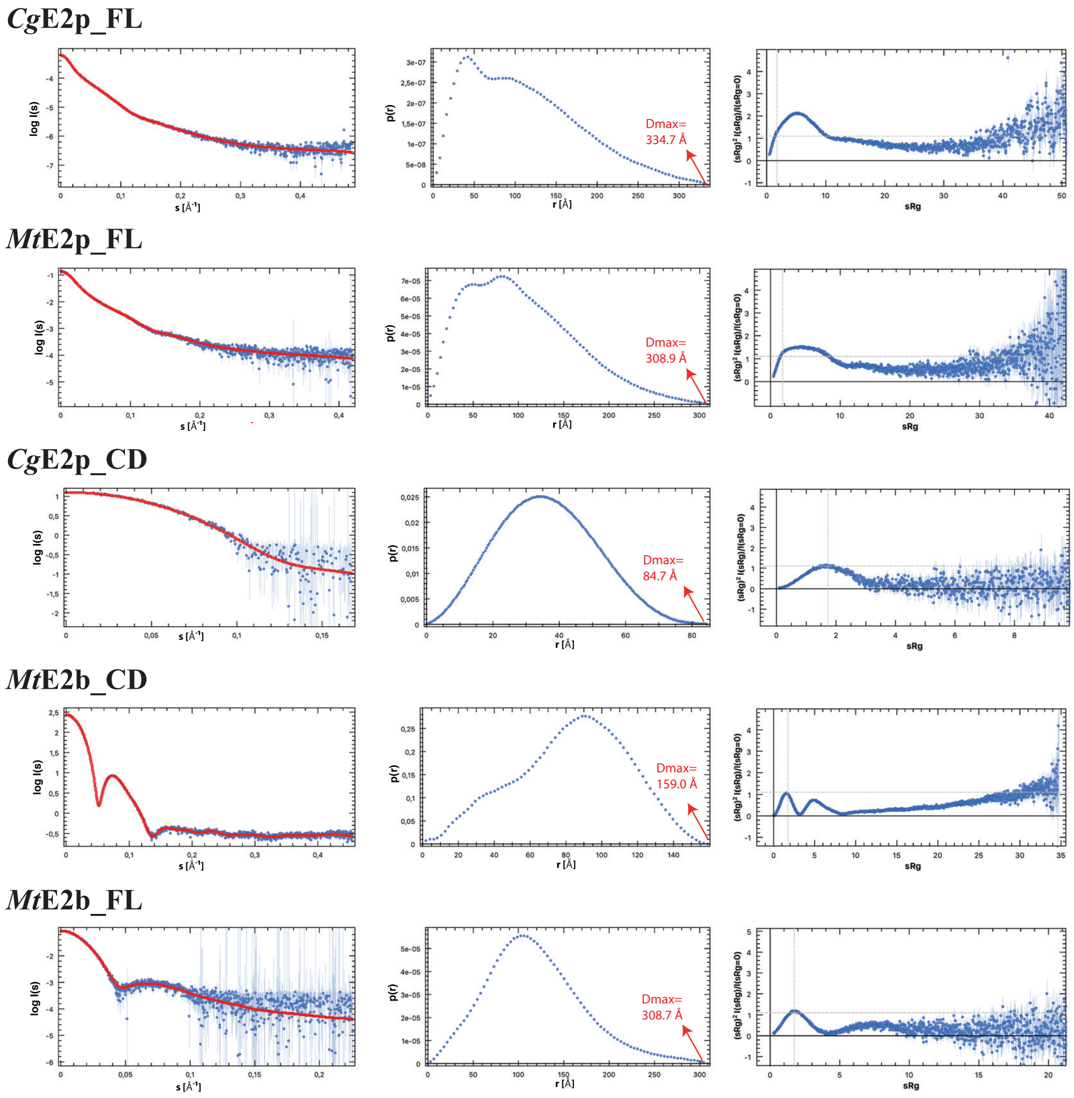


### Supplementary figure 1: SAXS primary data analysis.

The Pair distance distribution analysis was performed for each protein. Left: experimental scattering intensity (blue) and corresponding fitting curve (red). The central graph correspond to the Pair distance distribution function showing the estimated Dmax values. Right: dimensionless Kratky plot. Dashed lines indicate the position were a globular protein maximum is predicted to be located ($sRg=\sqrt{3}$ and $(sRg)^2I(s)/I(0) = 1.104$)[1,2].

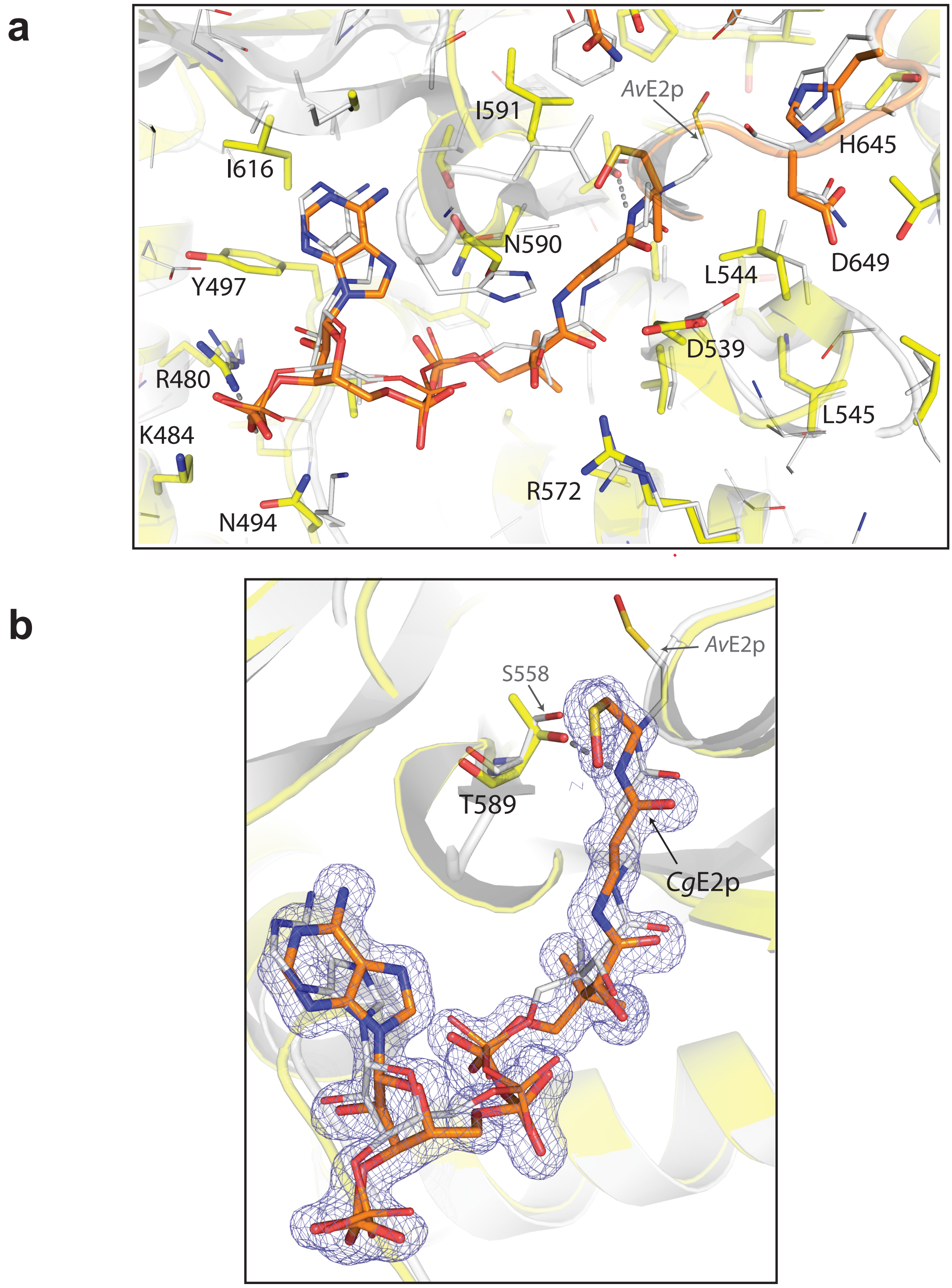


**Supplementary figure 2: Comparison of the CoA conformation within the active site of *Cg*E2p_CD and *Av*E2p_CD.**

**a**- Superposition of *Cg*E2p (yellow) and *Av*E2p (gray) active sites. Residue numbering corresponds to *Cg*E2p. The CoA Ligand is visible in the center (orange for *Cg*E2p and gray for *Av*E2p).
**b**- Zoomed region corresponding to the CoA binding site, showing the ligand density for *Cg*E2p (2Fo-Fc countured at the 1σ level, in blue). The terminal thiol group of CoA is found in a non-equivalent position to *Av*E2p_CD, and rather points to the opposite direction due to a reorientation of the cysteamine moiety close to 180° around N4P. Furthermore, the hydroxyl group of Thr589 (equivalent to Ser558 in *Av*E2p_CD) can form a H bond with the N4P atom from CoA to stabilize the negatively charged tetrahedral intermediate, and Fourier difference electron density maps are compatible with the presence of an oxygen atom at covalent binding distance (1.7 Å) from the CoA sulfur, suggesting the presence of, at least, a partial oxidation of the CoA thiol group to sulfenic acid.

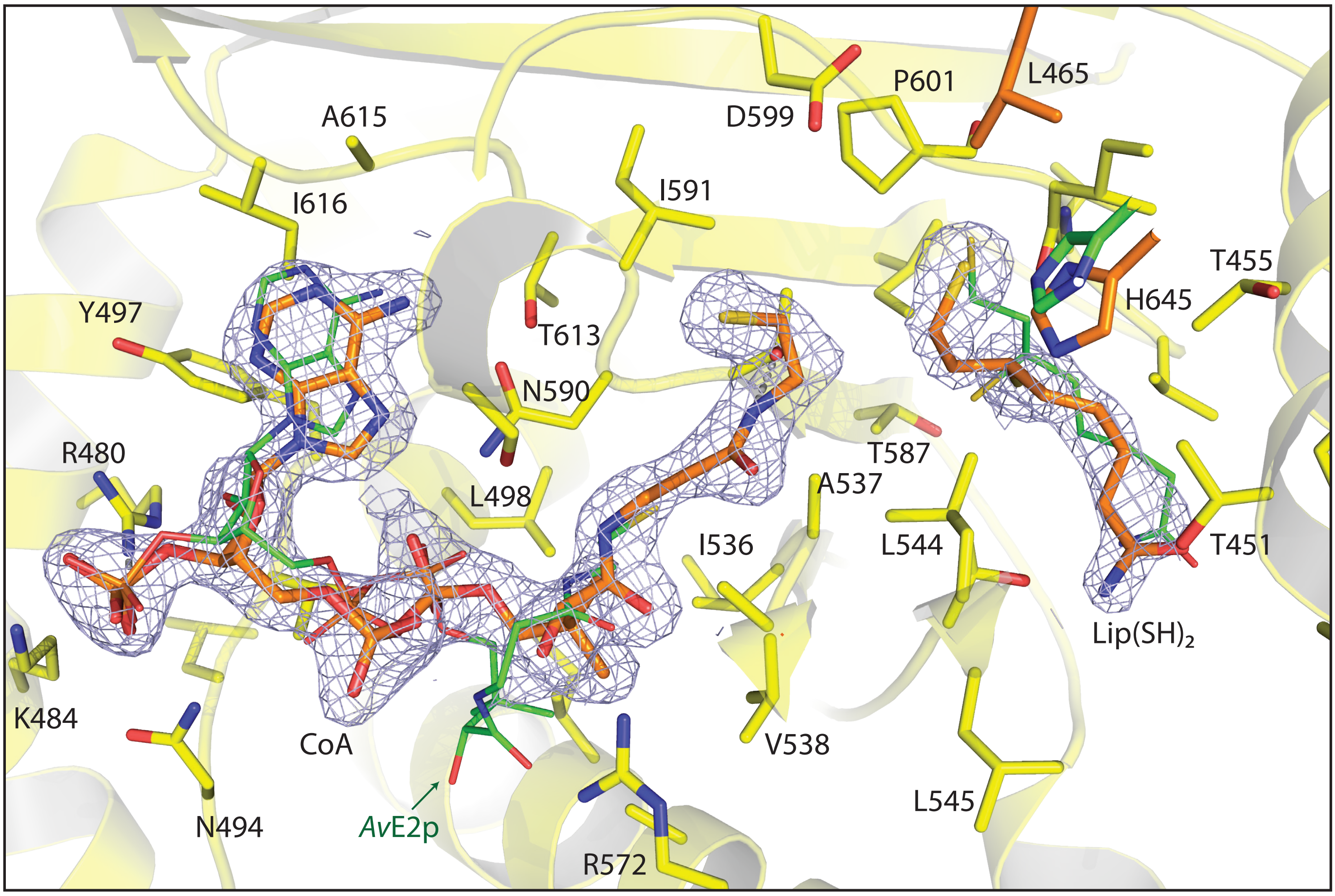


### Supplementary figure 3: Comparison of the ternary complex from *Cg*E2p_CD and *Av*E2p_CD.

Superposition of the ternary complex obtained for *Cg*E2p and *Av*E2p with its substrates CoA and Lipoamide. For *Cg*E2p residues and backbone are shown in yellow, side chains of active site residues are shown as sticks (and numbered) and both ligands are depicted as orange sticks. For *Av*E2p ligands are display in green (the *Av*E2p enzyme itself is not shown for clarity).
For *Av*E2p_CD, the ternary complex with CoA and Lip(SH)$_2$ presented CoA with an abortive, 'OUT' conformation in which the pantetheine chain did not reach the active site cleft but rather formed a left-handed helix with a series of intramolecular H-bonds (PDB entry 1EAB[3]). In the corynebacterial enzyme, despite a not strictly equivalent pose, Lip(SH)$_2$ makes active site interactions analogous to the ones reported in the case of *Av*E2p_CD, most notably the H-bond between the reactive sulphur (S8) and His645' provided by the opposite monomer, while the other sulphur atom (S6) is engaged in an H-bond with the carbonyl oxygen of Ile602, another conserved residue. Finally, the amide moiety is stabilized by H-bonds involving Thr451, the carbonyl group of Leu544 and the hydroxyl of a Ser residue preceding Leu437, left over from TEV digestion of the N-terminal His$_6$ affinity tag and forming crystal packing interactions. The other *Cg*E2p_CD monomer in the asymmetric unit, which forms a separate homotrimer via a crystallographic 3-fold axis, does not show any dihydrolipoamide bound, but a CoA molecule in the same orientation as the other protomer, and forming exactly the same interactions described before.

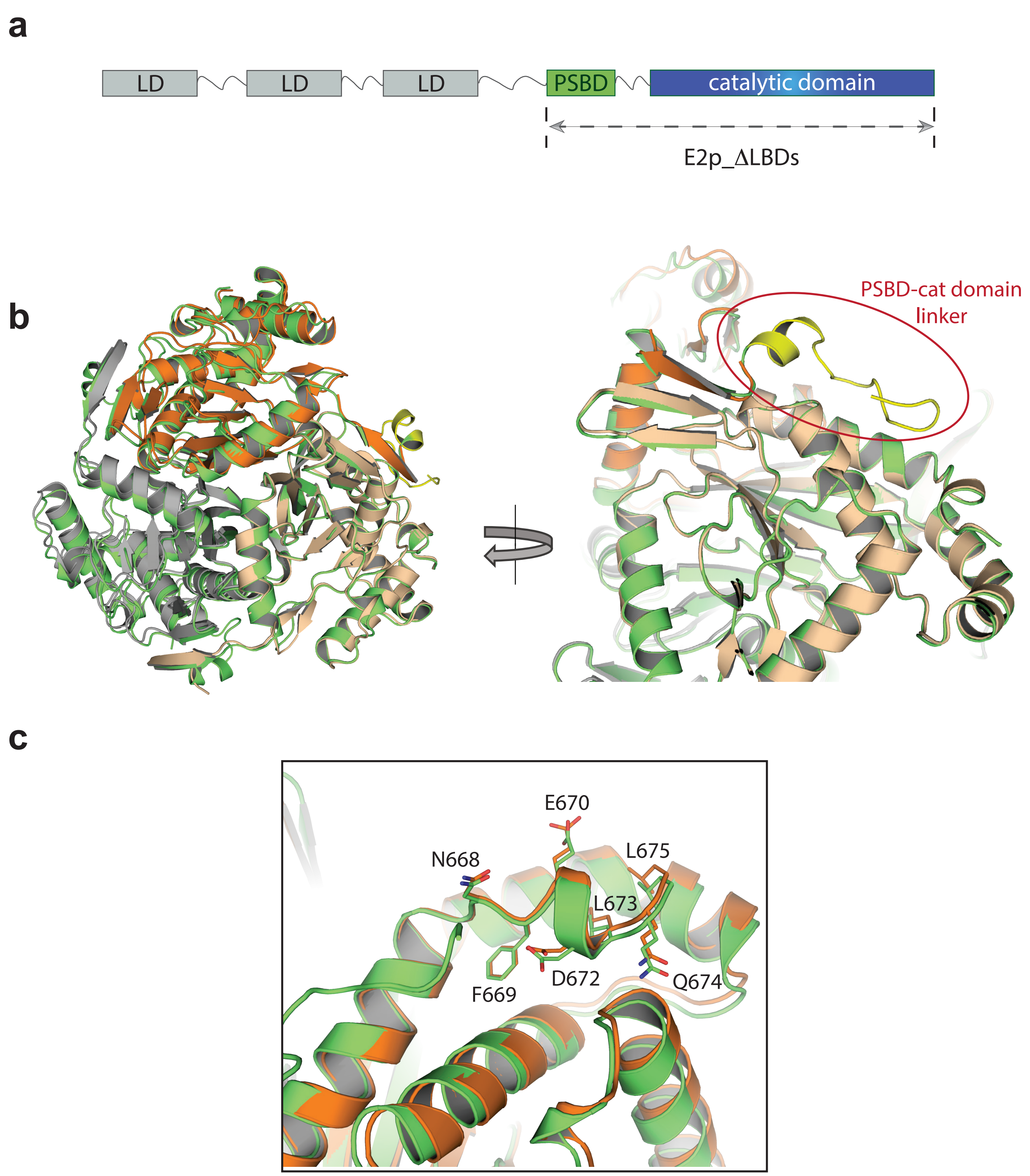


**Supplementary figure 4: Crystal structure of *Cg*E2p_ΔLBDs**

**a**- Schematic representation of the domain architechture of *Cg*E2p_FL. The arrow indicates the region selected for *Cg*E2p_ΔLBDs construction. **b**- Front and sideview of the superposition of the structures from *Cg*E2p_CD and *Cg*E2p_ΔLBDs (in orange and green respectively) showing the PSBD-catalytic domain linker present in the latter structure, in yellow. **c**- Zoomed region showing that the C-terminal α-helix, or TTI region, has the same conformation in both structures.

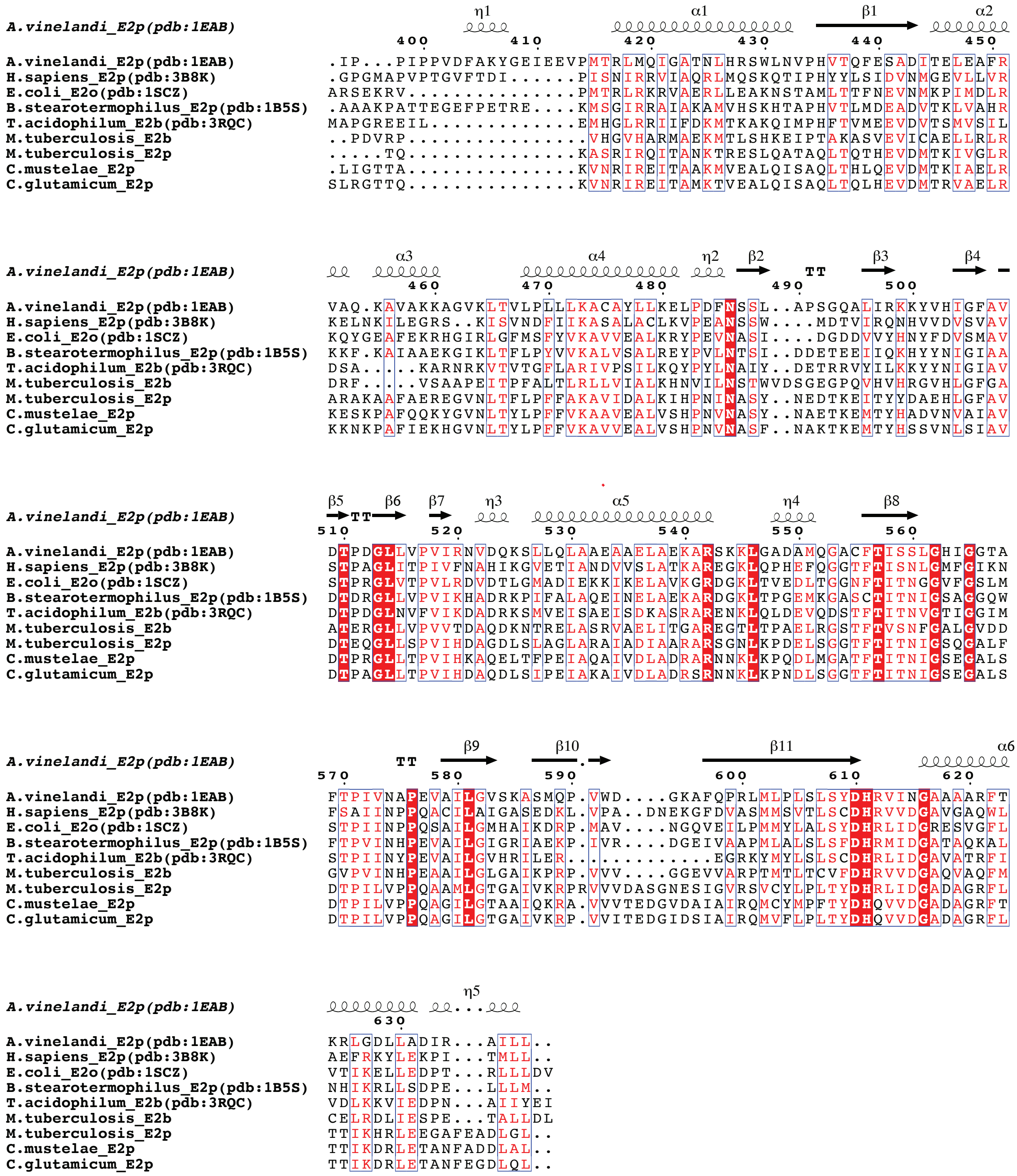


### Supplementary figure 5: sequence alignment of E2 proteins with known structure and *Mt*E2p, *Mt*E2b, *Cg*E2p and *Cm*E2p.

Sequence alignment including the catalytic domains of several E2 enzymes of known structure (namely, pdb entries: 1EAB, 3B8K, 1SCZ, 1B5S and 3RQC) together with *Mt*E2b, *Mt*E2p, *Cg*E2p and *Cm*E2p. As a reference, secondary structure features and residue number of *Av*E2p were superposed using Esprit3 (http://espript.ibcp.fr; [4]).

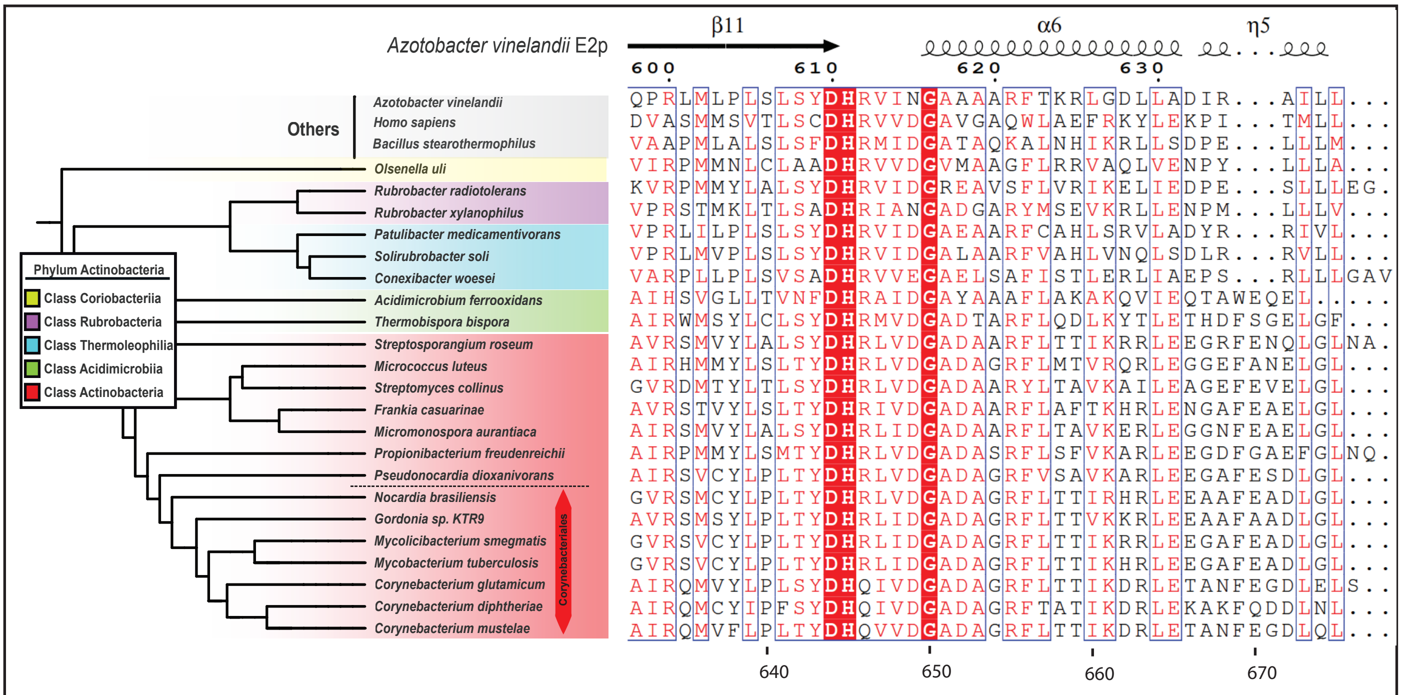


**Supplementary figure 6: Analysis of the distribution of the PCI inside Actinobacteria.**

Sequence alignment of several E2 of known structure toguether with E2 from organisms from the phylum Actinobacteria.Representative sequences from the phylum were selected to represent the 5 classes (Coriobacteriia in yellow, Rubrobacteria in violet, Thermoleophilia in cyan, Acidimicrobiia in yellow and Actinobacteria in red). Within the biggest class (class Actinobacteria or Actinomycetales) sequences selected represent different orders including the order Corynebacteriales, that include *M. tuberculosis* and *C. glutamicum*. The distribution of the selected genomes within the phylum can be observed in Figure 5B. Finally, the sequences were aligned using Muscle[5] and the presence of the PCI was analyzed in the region corresponding to the insertion observed in *Cg*E2p and *Mt*E2p. As a guide secondary structure features and residue number of *Av*E2p were superposed using ESPript3 (http://espript.ibcp.fr; [4]).

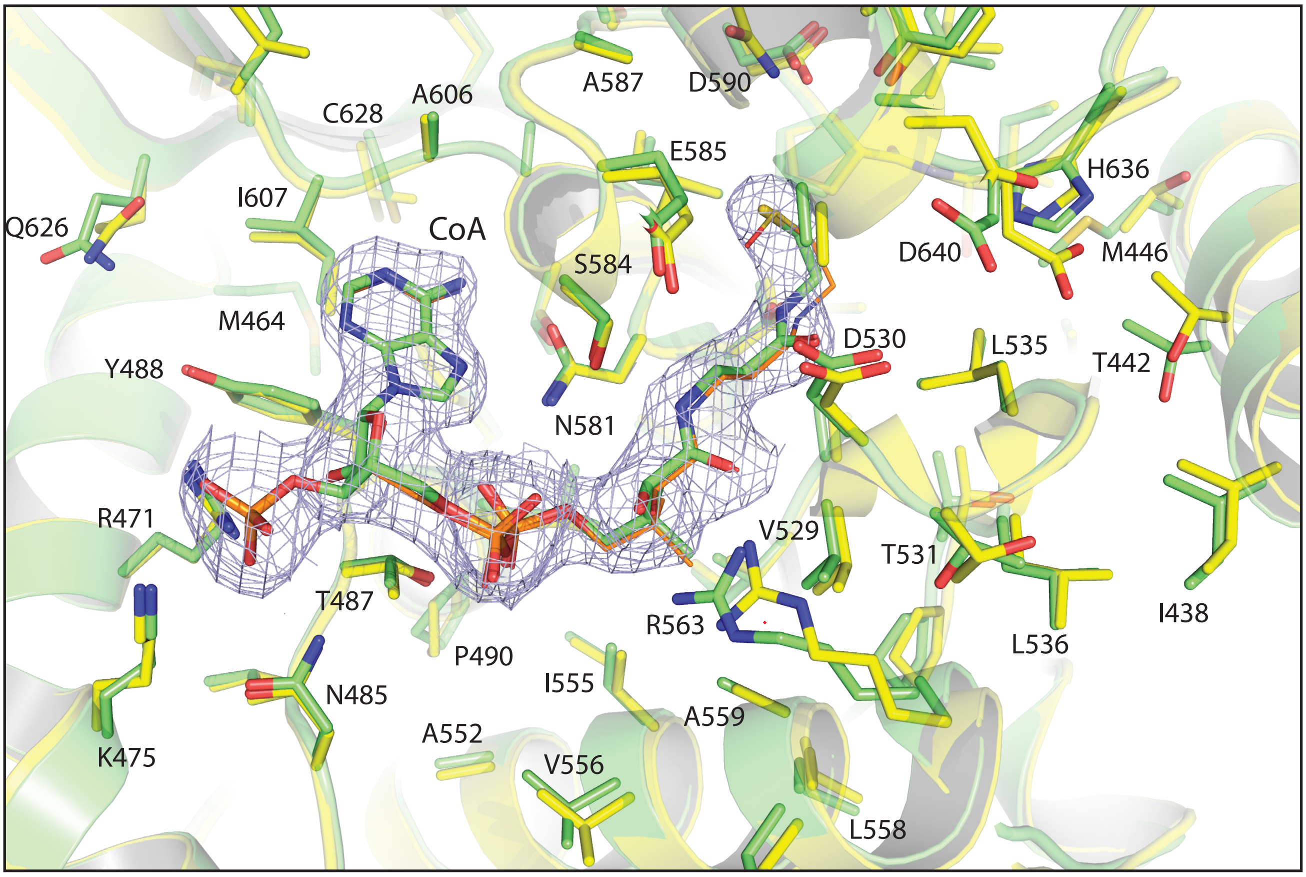


**Supplementary figure 7: Comparison of the active site pockets between *Cm*E2p_CD and *Cg*E2p_CD.**

Ribbon representation of *Cm*E2p_CD, in complex with CoA (green), superimposed to *Cg*E2p_CD (yellow) in complex with oxidised CoA (thinner sticks, orange). Side chains of residues within 10 Å from the ligand are shown in sticks; numerotation refers to *Cm*E2p_CD. The bound CoA molecule shows the same 'IN' conformation observed in *Cg*E2p_CD, including the peculiar, bent conformation of the terminal thiol group.

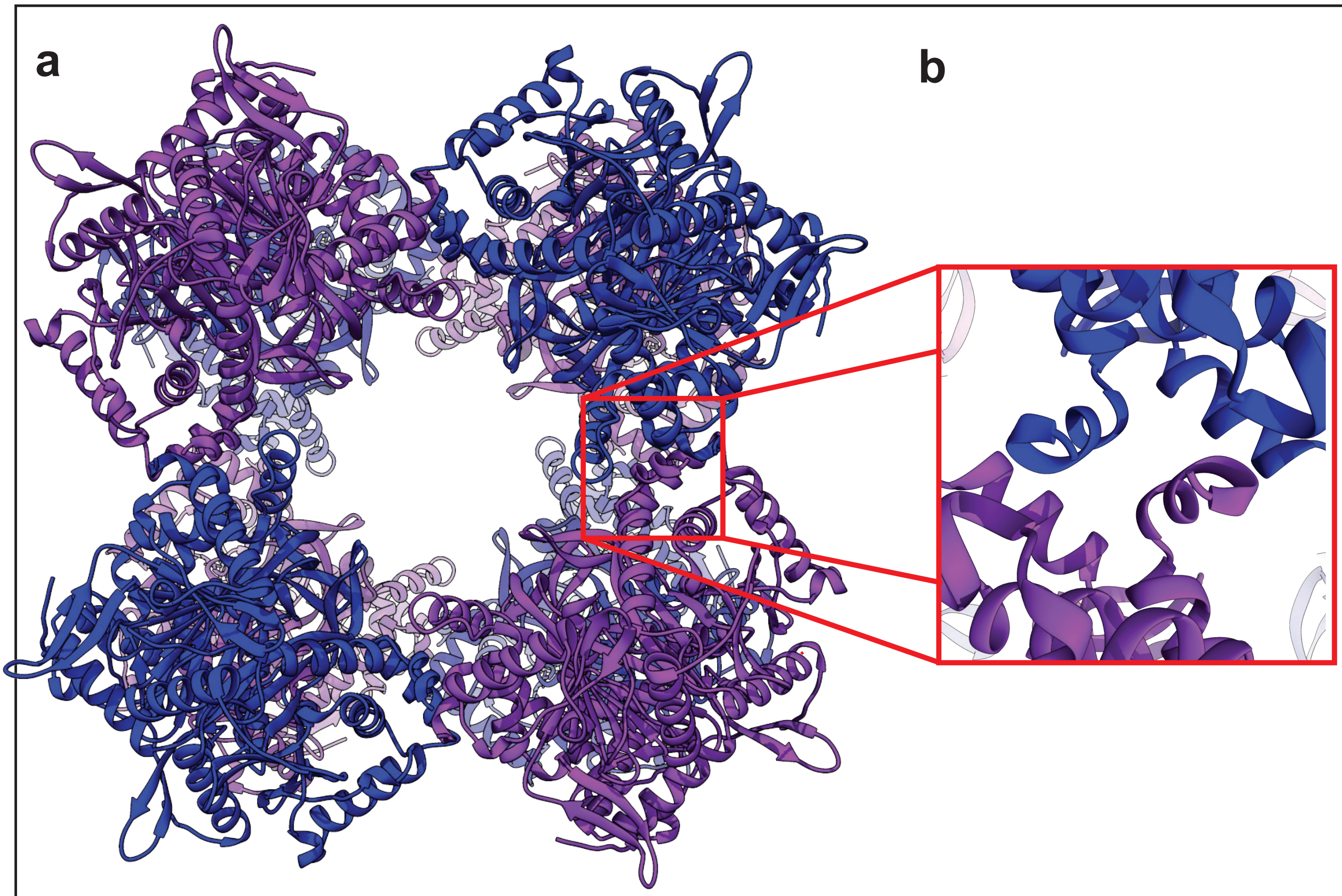


**Supplementary figure 8: Crystal structure of *Mt*E2b_CD.**

**a**- Ribbon representation of the crystal structure of *Mt*E2b_CD, which forms a 24-mer cubic structure entirely by crystallographic symmetry (the content of the asymmetric unit corresponds to a single monomer). Eight trimers are positioned at the corners of the cube (alternating blue and violet colours). **b**- Zoomed view of the region corresponding to the inter-trimer interaction where the role of the TTI-helix, main determinant of the trimer-trimer interaction, is highligted.

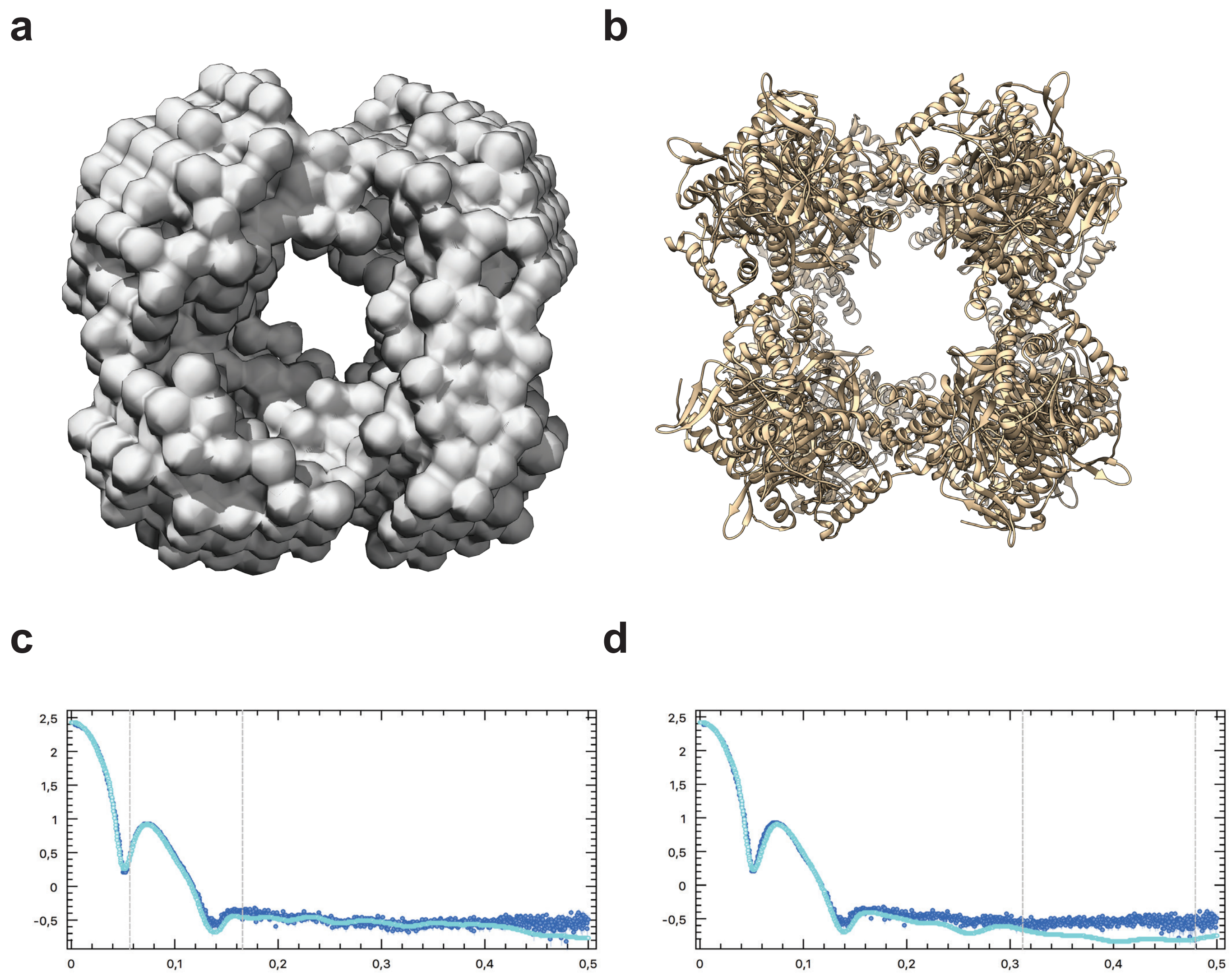


**Supplementary figure 9: *Mt*E2b_CD SAXS analysis.**

**a**- Front view from the DAMMIN generated *ab-initio* model of *Mt*E2b_CD reconstructed from the SAXS experimental data. **b**-Crystallographyc structure of the same protein (this work). **c**- Experimental scattering intensity (blue dots) plotted and compared with the theoretical fit obtained for the DAMMIN model (solid cyan line) **d**-Experimental and calculated scattering intensity for the crystall structure obtained using CRYSOL [6].

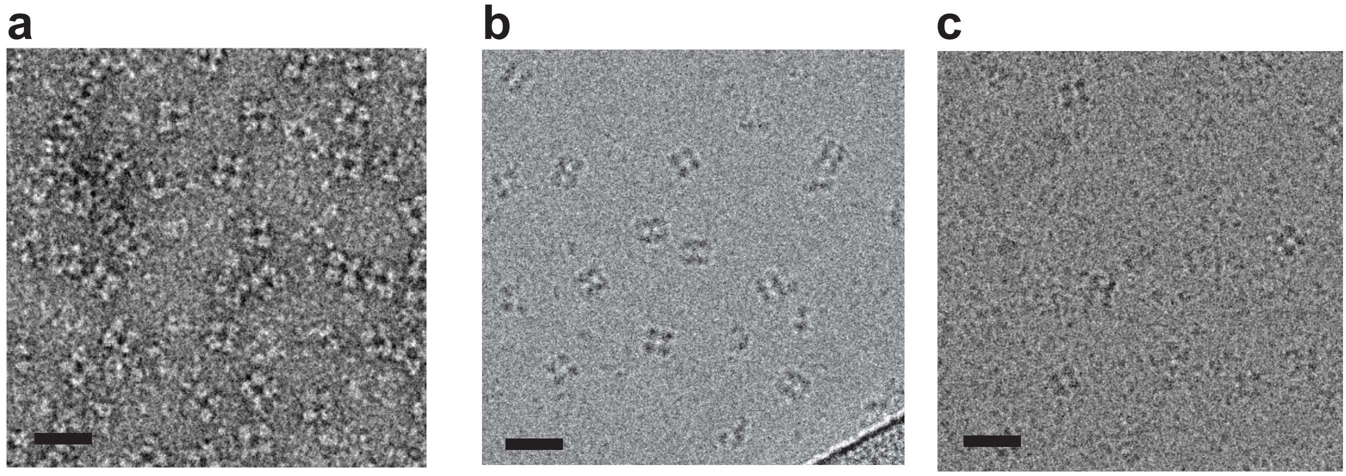


**Supplementary figure 10: Individual particles of *Mt*E2b visualized by EM.**
**a**- EM grid prepared using uranyl acetate as staining agent, showing the cubic particles of *Mt*E2b_CD. **b,c** - EM grid prepared in cryo condition showing cubic particles of *Mt*E2b_CD (**b**) and *Mt*E2b_FL (**c**). The scale bar represents 20 nm in all cases.

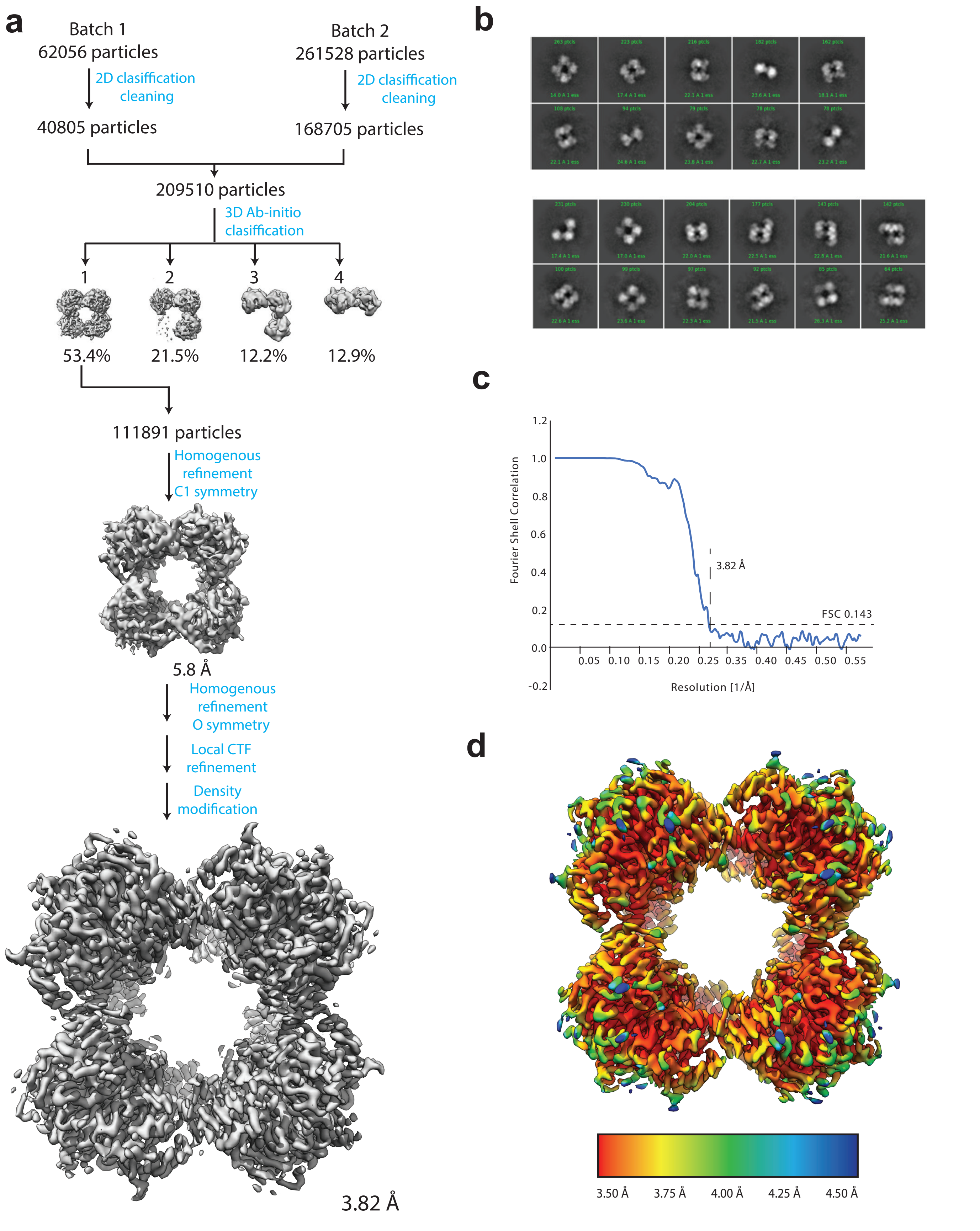


**Supplementary figure 11: CryoEM data analysis**

**a**- Flow chart of the CryoEM data processing from 2D classification to the final model. **b**- Representative 2D classes correponding to model 4 (top) and models 2 and 3 (bottom). **c**- FSC plots and resolution estimation using the gold-standard 0.143 criterion. **d**- Final refined map colored according to local resolution calculated with Blocres in Cryosparc [7,8,9].

Supplementary Table 1 | SAXS data collection and derived parameters

---

### Data collection parameters

| | |
|---|---|
| Instrument: | Beamline SWING (synchrotron SOLEIL) |
| Detector: | CCD-based AVIEX |
| Beam geometry: | 0.8 mm x 0.15 mm |
| Wavelength (Å): | 1.033 |
| s range ($Å^{-1}$)[a] : | 0.0064 < s < 0.50 |
| Exposure time (s): | 1.5 |
| Temperature (K): | 288 |

**Structural parameters**

| Protein | CgE2p_CD | CgE2p_FL | MtE2p_FL | MtE2b_CD | MtE2b_FL |
|---|---|---|---|---|---|
| $R_g$ Guinier [Å] | 27.7 ± 3.8 | 88.9 ± 2.53 | 80.97±1.06 | 62.11±1.85 | 92.79 ± 1.16 |
| $I_{(o)}$ Guinier [arbitraty units] | 12.38 ± 0.58 | 59 ± 2.1 | 0.13±0.006 | 260.94±7.2 | 0.086± 0.0068 |
| $R_g$ *P*(r) [Å] | 27.7 | 92.95 | 84.37 | 62.81 | 91.58 |
| $I_{(o)}$ *P*(r) [arbitraty units] | 12.38 | 59 | 0.13 | 267 | 0.08 |
| $D_{max}$ [Å] | 84.7 | 334.74 | 308.91 | 159,04 | 308,72 |
| Vol POROD $Å^3$ | 117250 | 578000 | 577000 | 1170000 | 3930000 |
| Mw Bayesian (kDa) | 76.3 | 242.6 | 318.4 | 479.1 | 1178.4 |
| MW credibility interval Bayesian | 71.4-81.9 | 221.0-372.7 | 221.0-372.7 | 264.2-556.0 | 916.6-inf |

---

[a] Momentum transfer |s| = 4πsin(θ)/λ

Abbreviations:

$M_W$: Molecular mass

$R_g$ : Radius of gyration

$D_{max}$ : Maximal particle dimension
